# Dual CRALBP isoforms unveiled: iPSC-derived retinal modelling and AAV2/5-RLBP1 gene transfer raise considerations for effective therapy

**DOI:** 10.1101/2024.04.24.590768

**Authors:** Krishna Damodar, Gregor Dubois, Laurent Guillou, Daria Mamaeva, Marie Pequignot, Nejla Erkilic, Carla Sanjurjo-Soriano, Hassan Boukhaddaoui, Florence Bernex, Béatrice Bocquet, Jerome Vialaret, Yvan Arsenijevic, T. Michael Redmond, Christopher Hirtz, Isabelle Meunier, Philippe Brabet, Vasiliki Kalatzis

## Abstract

Inherited retinal diseases (IRDs) are clinically and genetically heterogeneous disorders characterised by progressive vision loss. Over 270 causative genes have been identified and variants within the same gene can give rise to clinically distinct disorders. Human induced pluripotent stem cells (iPSCs) have revolutionised disease modelling, by allowing pathophysiological and therapeutic studies in the patient and tissue context. The IRD gene *RLBP1* encodes CRALBP, an actor of the rod and cone visual cycles in the retinal pigment epithelium (RPE) and Müller cells, respectively. Variants in *RLBP1* lead to three clinical subtypes: Bothnia dystrophy, Retinitis punctata albescens and Newfoundland rod-cone dystrophy. We modelled *RLBP1*-IRD subtypes by patient-specific iPSC-derived RPE and identified pertinent therapeutic read-outs. We developed an AAV2/5-mediated gene replacement strategy and provided a proof-of-concept in the *ex vivo* human models that was validated in an *in vivo Rlbp1^−/−^* murine model. Most importantly, we identified a previously unsuspected smaller CRALBP isoform that is naturally and differentially expressed in both human and murine retina. The new isoform arises from an alternative methionine initiation site and plays a role in the visual cycle. This work provides novel insights into CRALBP expression and *RLBP1*-associated pathophysiology and raises important considerations for successful gene supplementation therapy.

## Introduction

Light absorption by the retina initiates a series of chemical and electrical signals that are transmitted to the brain to create visual perception (1). The light-sensitive retinal cells are the rod and cone photoreceptors. In the human retina, the rods are preferentially located peripherally and are active in dim light, while the cones are preferentially located centrally and are active in bright light. Inherited retinal dystrophies (IRDs) are a group of disorders characterised by photoreceptor degeneration and a progressive loss of vision (2). They are clinically heterogenous with peripheral (rod loss) or central (cone loss) vision loss, and variable ages of onset and severity. Although monogenic, IRDs are linked to variants in over 270 genes important for the photoreceptors and/or the underlying support tissue, the retinal pigment epithelium (RPE). Furthermore, variants in the same gene can give rise to clinically distinct disorders.

One such example is *RLBP1*. *RLBP1* variants are associated with the autosomal recessive IRD, retinitis punctata albescens (RPA; OMIM 136880) (3–5). RPA is characterised by a childhood onset of night blindness, elevated dark adaptation thresholds and uniform white dots across the fundus. The disease progresses to a generalised atrophy of the retina and macula (cone-rich centre), leading to severe visual impairment from 40 years of age. *RLBP1* variants are also associated with two more severe clinical subtypes (6): Swedish Bothnia dystrophy (BD; OMIM 607475) that is characterised by an early involvement of the macula (7, 8), and Canadian Newfoundland Rod-Cone Dystrophy (NFRCD; OMIM 607476) that is characterised by night blindness from infancy and severe vision loss from 20 years of age (9). *RLBP1* variants have also been associated with a stationary form of night blindness, Fundus albipunctatus (OMIM 136880), but which likely represents a mis-diagnosis of slowly progressive RPA in young patients (10).

*RLBP1* encodes the cellular retinaldehyde binding protein, CRALBP (11). CRALBP is an actor of the visual cycle localised between the photoreceptor outer segments (POS) and the RPE to regenerate the visual chromophore 11-*cis* retinal following light absorption (12). The cycle involves a number of key steps: following photoisomerization of 11 *cis*-retinal and its reduction in the POS to all-*trans* retinol, the latter passes into the RPE and is esterified to all-*trans* retinyl esters by lecithin:retinol acyltransferase (LRAT); all-*trans* retinyl esters are then isomerised to 11-*cis* retinol by RPE65; 11-*cis* retinol is oxidised to light-sensitive 11-*cis* retinal by the retinal dehydrogenases (RDH). The rate-limiting step of the visual cycle is the isomerisation by RPE65 (13–15), which is accelerated by CRALBP (16). CRALBP chaperones 11-*cis* retinol to the RDH enzymes for oxidation and then the 11-*cis* retinal to the RPE membrane for transfer to the POS by the interphotoreceptor retinal binding protein (IRBP). The resulting retinoid depletion in the RPE by CRALBP stimulates the activity of RPE65. It is generally considered that there is a separate visual cycle for rods and cones (17). Rods use the canonical RPE cycle whereas the cone visual cycle is thought to be mediated by the Müller cells, the glial cells of the retinal, to specifically enable cones to quickly recover from bright light exposure (18). CRALBP has been shown to cooperate with retinal G-protein-coupled receptor (RGR) opsin, a photoisomerase, like CRALBP found in both the RPE and Müller glia, in a photic visual cycle based in both tissues and supplying both central and peripheral retina (19). CRALBP acts to selectively capture 11-*cis* retinal produced by RGR. In addition, CRALBP co-precipitates with another potential cone isomerase, dihydroceramide desaturase-1 in Müller cells (20). Taken together, CRALBP is essential for visual cycle kinetics in both rod- and cone-mediated vision (21).

Currently no universal treatment exists for IRDs, but a number have benefited from gene supplementation clinical trials (22). The treatment of IRDs due to a visual cycle defect became a reality when Luxturna, an adenovirus-associated virus serotype 2 (AAV2/2) vector carrying *RPE65* under the control of the ubiquitous CAG promoter, was market-approved (23). RPE65 deficiency causes all-*trans* retinyl ester accumulation leading to delayed or no 11-*cis* retinal production (24). This is quite similar to the effect of CRALBP deficiency (16), but more severe. Thus, considering the Luxturna success story, RPA represented an excellent candidate for gene therapy. Along this line, in 2015, Choi *et al.* reported that a self-complementary (sc) AAV2/8 vector expressing *RLBP1* under a short version of the endogenous human promoter (hRLBP1), improved the rate of dark adaptation in the *Rlbp1^−/−^* mouse model (25), and a clinical trial (NCT03374657) was initiated in 2018, the results of which are as yet unpublished.

In parallel, as we previously showed that AAV2/5 is the most effective serotype at transducing human RPE (26), we initiated a proof-of-concept study with an AAV2/5 vector expressing *RLBP1* under control of the CAG promoter. To be clinically relevant, we first modelled *RLBP1*-associated IRDs by generating induced pluripotent stem cell (iPSC)-derived RPE from patients carrying *RLBP1* variants associated with RPA, BD and NFRCD. We assayed the effect of these variants on CRALBP expression and RPE function to identify pertinent read-outs to assay therapeutic rescue. We then performed proof-of-concept studies for AAV2/5-mediated *RLBP1* supplementation *ex vivo* in the *RLBP1* iPSC-derived RPE and *in vivo* in the *Rlbp1^−/−^*mouse model. Interestingly and unexpectedly, this proof-of concept study, has brought to light novel findings regarding CRALBP. We show here that the functionally validated AAV2/5-CAG-RLBP1 vector, encodes two CRALBP isoforms, in contrast to the previously published AAV2/8-hRLBP1-RLBP1 vector (25). Moreover, we show that these isoforms are naturally and differentially expressed both in the mouse and human retina. Finally, we demonstrate that the smaller isoform arises from a second methionine codon and that it also plays a role in the visual cycle. Taken together, this work provides novel insights into CRALBP expression and *RLBP1*-associated pathophysiology and raises important considerations for successful gene supplementation therapy.

## Results

### Functionality of *RLBP1* iPSC-derived RPE correlates with clinical severity

To perform disease modelling of *RLBP1*-associated IRDs, and to generate a pertinent human model for proof-of-concept studies, we cultured skin fibroblasts from *RLBP1* patients presenting clinical subtypes (6): a 32 year-old woman (referred to as RPA1) with BD (Fig. 1A), a 14 year-old boy (RPA2) with classical RPA (Fig. 1B) and a 31 year-old man (RPA3) with severe NFRCD (Fig. 1C). We reprogrammed the fibroblasts into iPSCs, which were quality controlled for Sendai vector clearance by RT-PCR (Fig. S1A), pluripotency by qPCR (Fig. S1B), trilineage differentiation by teratoma assays (Fig. S1C), genetic integrity by karyotype analyses (Fig. S1D), and for the presence of the causative *RLBP1* variants. By PCR amplification and Sanger sequencing we confirmed the c.333T>G (p.Tyr111*) variant in exon 5 and c.700C>T (p.Arg234Trp) in exon 8 for RPA1 (Fig. 1D), and c.25C>T (p.Arg9Cys) in exon 4 and c.333T>G in exon 5 for RPA2 (Fig. 1E), and by long-range PCR analysis, we detected a 300-bp fragment corresponding to the homozygous exon 7 to 9 deletion (3) for RPA3 (Fig. 1F), compared to an ∼8-kb fragment for control iPSCs. The position of the variants on the *RLBP1* gene are schematized in Fig. 1G.

**Figure 1:**
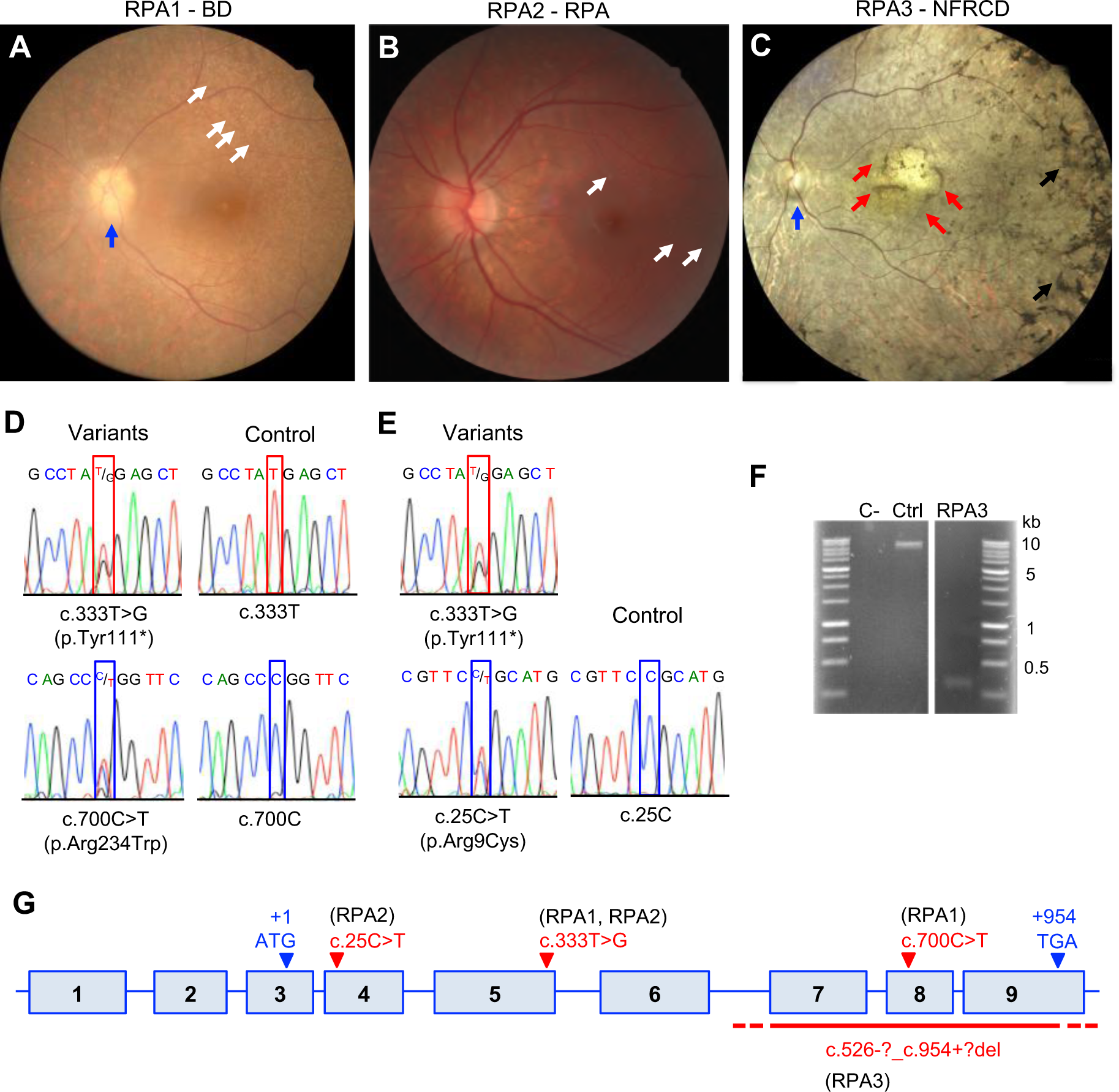
Clinical phenotype and genotype of *RLBP1* patients. Colour fundus imaging of individual: RPA1 at 28 years of age presenting with BD showing multiple white dots (white arrows), an irregular pale optic nerve head with papillary drusen (blue arrow) and reduced retinal vessel caliber (**A**); RPA2 at 23 years of age presenting with mild RPA showing multiple white dots (white arrows) but preserved papillary head and retinal vessels (**B**), and RPA3 presenting with severe NFRCD at 45 years of age showing nummular pigmentary changes (black arrows), a pale optic nerve head (blue arrow) and an altered and atrophic macula (red arrows) (**C**). Sanger sequencing of the *RLBP1* variants c.333T>G and c.700C>T (**D**) and c.25C>T and c.333T>G (**E**) in the iPSCs of individuals RPA1 and RPA2, respectively. **F**) Long-range PCR amplification of the region flanking the homozygous exon 7 to 9 deletion carried by RPA3. C-, water control. Ctrl, Control DNA. **G**) Schematic representation of *RLBP1* with the position of the variants carried by RPA1, RPA2 and RPA3 indicated in red; the position of the start and stop codons are indicated in black.

We then differentiated the iPSCs into RPE and by immunofluorescence (IF) studies we detected expression of characteristic markers in all patient lines: the apical adherens marker ZO1 (Fig. 2A), the basolateral chloride channel BEST1 (Fig. 2B), the perinuclear and cytosolic visual cycle markers LRAT (Fig. 2B) and RPE65 (Fig. 2C), respectively, and the apical microvilli marker MERTK (Fig. 2C); representative data shown for the severe RPA3 line. Transepithelial resistance (TER) recordings increased weekly for the control and patient iPSC-derived RPE to values >150 Ω/cm^2^ (Fig. 2D), indicative of a tight monolayer (27). We assayed the general functionality of the iPSC-derived RPE by feeding with FITC-labelled photoreceptor outer segments (POS) and quantifying the percentage of POS-containing cells (Fig. 2E) and their mean intensity of fluorescence (Fig. 2F) by flow cytometry. Overall, this characteristic function was not significantly different between control, RPA1 and RPA2 RPE, whereas RPA3 RPE showed significantly reduced phagocytosis compared to the other lines.

**Figure 2:**
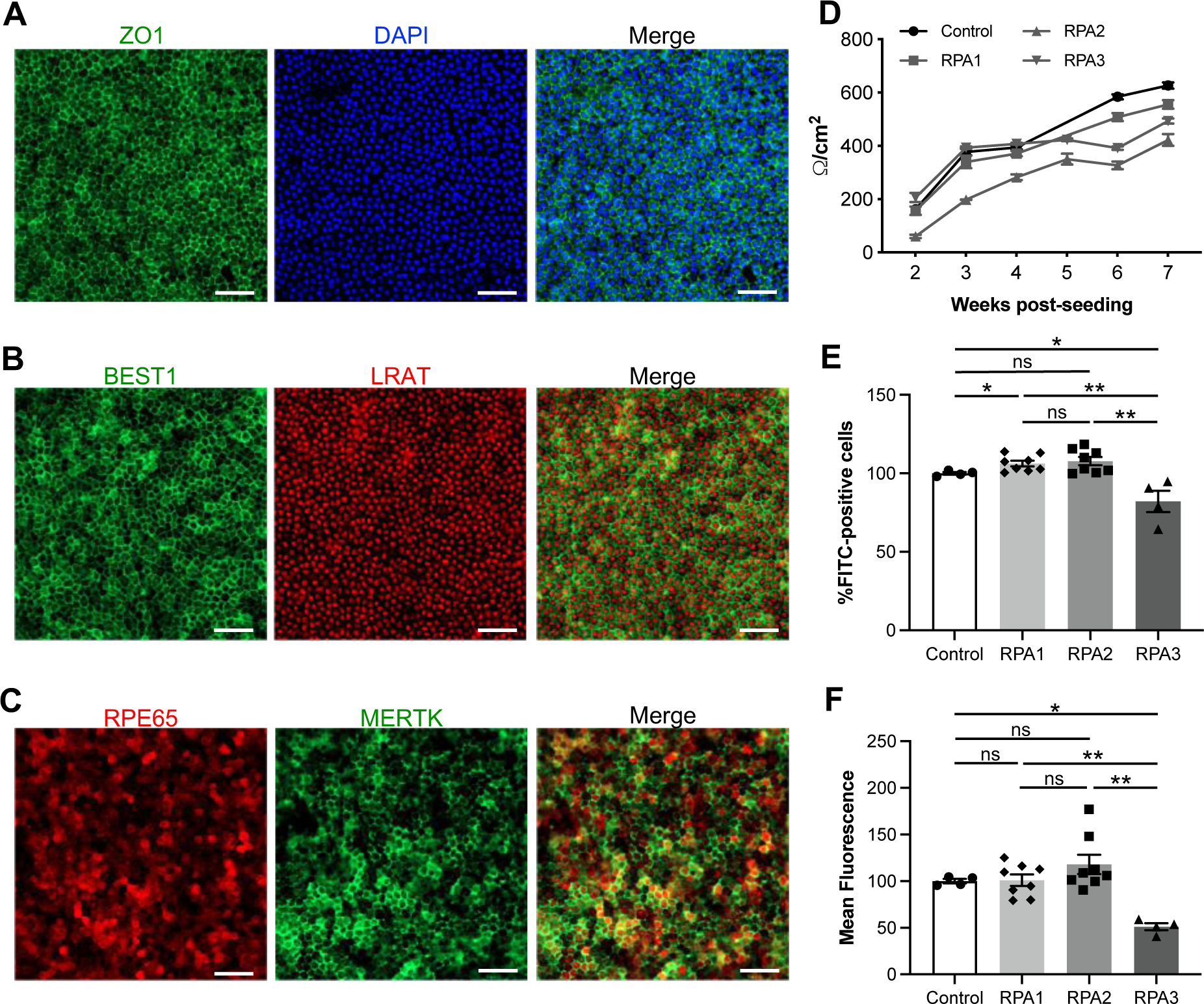
Morphological and functional characteristics of patient iPSC-derived RPE. Representative IF images of the iPSC-derived RPE from patient RPA3 stained for the markers ZO1 (**A**), BEST1 and LRAT (**B**), and RPE65 and MERTK (**C**); nuclei are stained in blue. Scale bars = 50 µm. **D**) Weekly TER measurements of control, RPA1, RPA2 and RPA3 iPSC-derived RPE expressed in ohms/cm2. Data are represented as mean ± SEM of recordings from 5 inserts (control and RPA1) or 6 inserts (RPA2 and RPA3). Flow cytometry analysis of the percentage of iPSC-derived RPE cells that phagocytosed FITC-labelled POS (**E**) and the mean fluorescence intensity following phagocytosis (**F**) expressed relative to control. Data are represented as mean ± SEM; \**p*<0.05; \*\**p*<0.01; ns, non-significant; *n* is indicated by the symbols.

Taken together, iPSC-derived RPE from *RLBP1* patients presents as a morphologically characteristic monolayer with altered functionality for the most severe clinical phenotype.

### CRALBP expression in iPSC-derived RPE depends on the causative *RLBP1* variant

We next assayed *RLBP1* expression by qPCR and it was detected in the control iPSC-derived RPE and not in undifferentiated iPSCs (Fig. 3A). We detected significantly reduced *RLBP1* levels in the iPSC-derived RPE lines RPA1 (0.2-fold of control) and RPA2 (0.5-fold), which carried a combination of missense and nonsense alleles, and RPA3 (0.13-fold), which carried the large exon 7 to 9 deletion, compared to controls. This expression profile was confirmed using a second set of primers in different exons (Fig. S2A). Consistent with varying mRNA levels between patients, western blot analysis of iPSC-derived RPE cell lysates detected CRALBP in RPA2 RPE, at 0.2-fold of control levels, but not in RPA3 RPE (Fig. 3B). Surprisingly, CRALBP expression was not detected in RPA1 RPE.

**Figure 3:**
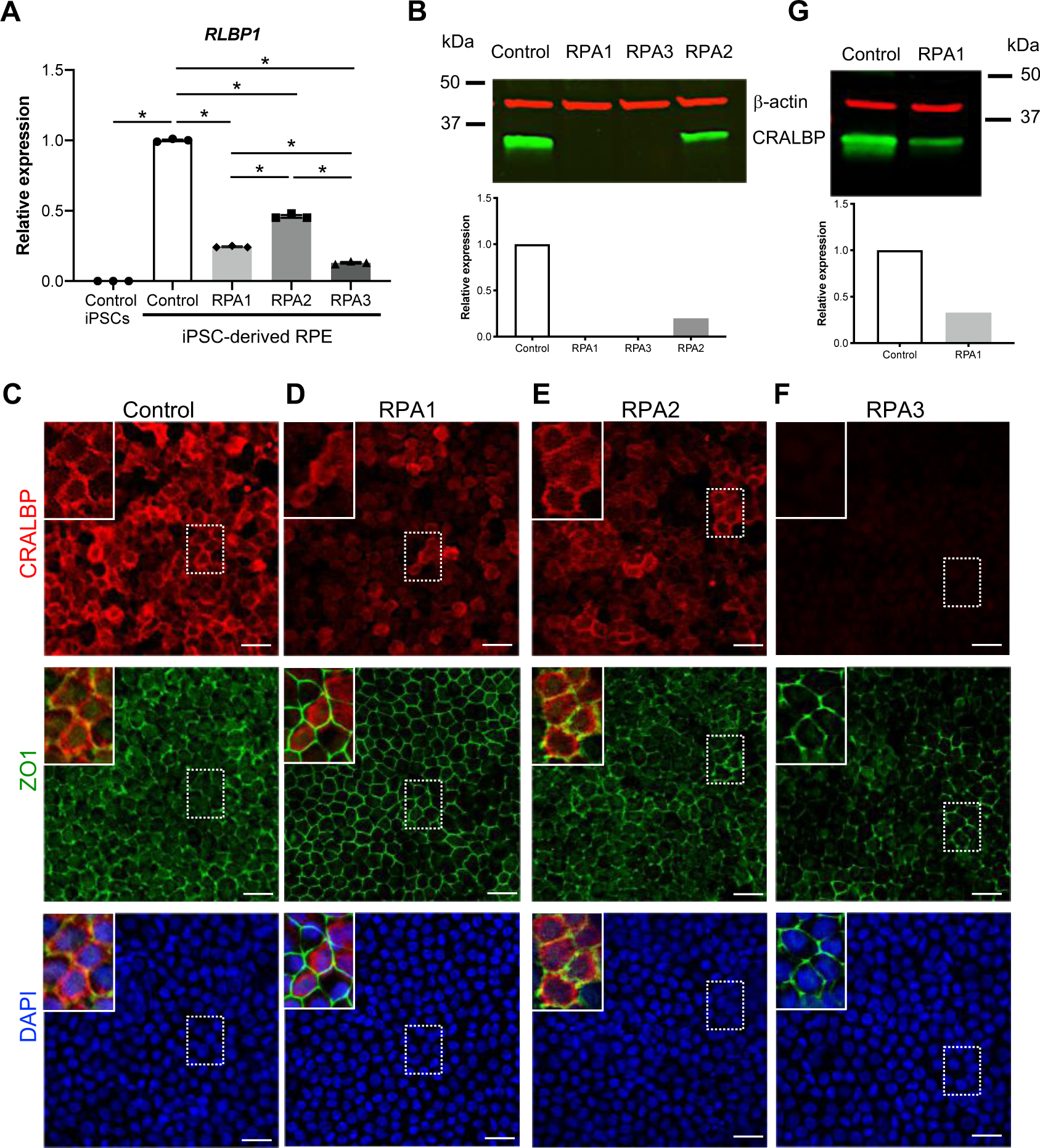
*RLBP1* and CRALBP expression in iPSC-derived RPE. **A**) qPCR analysis of *RLBP1* expression using exon 6 primers in control iPSCs, and control and patient iPSC-derived RPE. Data are represented as mean ± SEM and expressed relative to control; \**p*<0.05; *n* is indicated by the symbols. **B**) Western blot analysis of CRALBP expression using a rabbit monoclonal antibody (green) in 10 µg of control and patient iPSC-derived RPE cell lysates. The graphs represent the quantification of CRALBP relative to the loading control, β-actin (red). Representative IF images of control (**C**), RPA1 (**D**), RPA2 (**E**) and RPA3 (**F**) iPSC-derived RPE stained for CRALBP and ZO1 and shown as individual channels; nuclei are labelled in blue. Scale bars = 20 µm. Insets, 2-fold magnification of the boxed areas for each panel shown as merged images with the preceding channel (top row - red alone, middle row - red and green), bottom panel (red, green and blue). **G**) Western blot analysis of CRALBP expression in control and RPA1 iPSC-derived RPE directly resuspended in Laemmli buffer and migrated on a gradient 4-20% polyacrylamide gel. The graph represents the quantification of CRALBP relative to the loading control, β-actin.

We then performed IF studies to further verify CRALBP expression. In control iPSC-derived RPE, CRALBP staining was diffused throughout the cytosol but highly concentrated on the inner side of the plasma membrane (Fig. 3C), consistent with our previous reports (28, 29). Unexpectedly, in RPA1 RPE, CRALBP expression was faint and partially mis-localised; the staining was mainly diffused in the cytosol and unevenly distributed near the plasma membrane (Fig. 3D). By contrast, CRALBP expression in RPA2 RPE showed a similar pattern to control RPE although less intense (Fig. 3E). Consistent with the western blot data, CRALBP was not detected for RPA3 (Fig. 3F). Due to the discrepancy in CRALBP detection for RPA1 between the western blot analysis of RPE cultured in 24-well plates, and the IF data for RPE cultured on transwell inserts, we performed additional western blot analyses on iPSC-derived RPE cultured on transwell inserts. Under these conditions, CRALBP was detectable, at levels 0.3-fold that of control (Fig. 3G).

In conclusion, the RPE from the BD and RPA patients has reduced *RLBP1* expression but detectable CRALBP, whereas the RPE from severe NFRCD has no detectable CRALBP.

### *RLBP1* iPSC-derived RPE shows retinoid accumulation

CRALBP is an 11-*cis*-retinoid-binding protein that drives the isomerisation reaction performed by RPE65 (30). We hence assayed retinoid accumulation in the RPE associated with the two most extreme clinical phenotypes, mild RPA (RPA2) and severe NFRCD (RPA3). To this end, we incubated iPSC-derived RPE with all-*trans* retinol to mimic the visual cycle and assayed all-*trans* retinyl ester levels by HPLC. The RPE of both patients showed significantly higher all-*trans*-retinyl ester levels compared to control RPE (Fig. 4A). Furthermore, in native RPE, retinoids accumulate in a specialised lipid droplet termed a retinosome (31), which can be assayed by transmission electron microscopy (TEM). We performed TEM analysis and observed a similar polarised (apical microvilli, basal nuclei) structure in both control and patient iPSC-derived RPE, with intracellular droplets readily visible in patient RPE (Fig. 4B). The number of droplets was significantly higher in RPA2 and RPA3 RPE compared to controls from 2- to 3-mo post-seeding. Furthermore, with prolonged culture (4- to 7-mo post-seeding), the number of droplets were 2.3-fold higher in the patient RPE, compared to the earlier time-point, and were still significantly more abundant than in control RPE (Fig. 4C).

**Figure 4:**
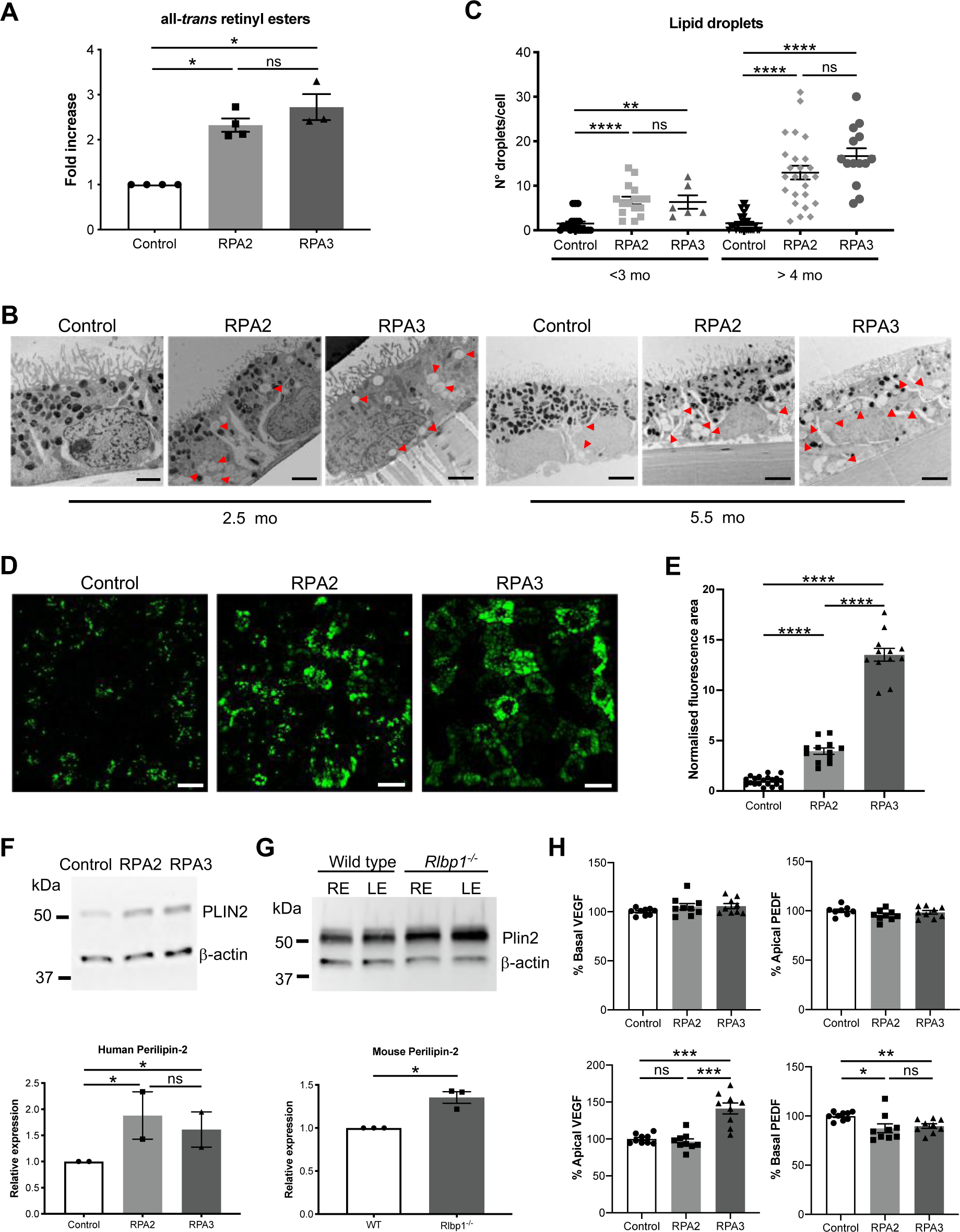
Retinosome accumulation in iPSC-derived RPE. **A**) HPLC of all-*trans* retinyl ester levels in control, RPA2 and RPA3 RPE expressed as a fold increase over control. Data are represented as mean ± SEM; *p<0.05; *n* is indicated by the symbols. **B**) Representative TEM images of 2.5- and 5.5-month-old control, RPA2 and RPA3 iPSC-derived RPE. Lipid droplets are indicated by red arrowheads. Scale bars = 2 µm. **C**) Quantitative analysis of the number of droplets/cell following TEM analysis of control and patient iPSC-derived RPE grouped according to age. Data are represented as mean ± SEM; \*\*\**p*<0.001; \*\*\*\**p*<0.0001; *n* is indicated by the symbols. **D**) Representative images of multiphoton microscopy of fluorescent retinosomes in 7-month-old control and patient iPSC-derived RPE. Scale bars = 15 µm. **E**) Quantitative analysis of the area of fluorescence normalized to the area of the image. Data are represented as mean ± SEM; \*\*\*\**p*<0.0001; *n* is indicated by the symbols. **F**) Representative western blot analysis of human perilipin-2 (PLIN2) expression in control, RPA2 and RPA3 iPSC-derived RPE; β-actin was the loading control. Graph represents the quantification of PLIN2 normalised to β-actin and expressed relative to control in 2 independent experiments. Data are represented as mean ± SEM; *p<0.05. **G**) Representative western blot analysis of murine perilipin-2 (Plin2) expression in the right (RE) and left (LE) eye of wild type and *Rlbp1^−/−^* RPE; β-actin was loading control. Graph represents the quantification of Plin2 normalised to β-actin and expressed relative to control in 3 independent experiments. Data are represented as mean ± SEM; \**p*<0.05. **G**) ELISA of VEGF and PEDF secreted apically and basally from iPSC-derived RPE. Data are represented as mean ± SEM and expressed as a percentage of control; *p<0.05; **p<0.01; ***p<0.001; ****p<0.0001; ns, non-significant; *n* is indicated by the symbols.

Reportedly, retinosomes can also be detected using multiphoton microscopy from wavelengths 720 nm to 780 nm (32). We thus assayed iPSC-derived RPE using multiphoton microscopy and readily detected fluorescent vesicles that appeared larger in RPA2 and RPA3 RPE, compared to control tissue (Fig. 4D). We quantified the area of fluorescence and confirmed a significant increase in the *RLBP1* RPE compared to control RPE (Fig. 4E). To definitively identify the lipid droplets as retinosomes, we tested expression of the specific marker, perilipin-2, by western blot analysis and detected significantly higher levels in RPA2 and RPA3, compared to control, RPE (Fig. 4F). As we had access to the *Rlbp1^−/−^* mouse model (see below), we further validated these data by western blot analysis of murine RPE and detected significantly higher perilipin 2 levels in *Rlbp1^−/−^,* compared to wild type, RPE (Fig. 4G). Lastly, we previously showed retinosome accumulation in association with defective polarised secretion in iPSC-derived RPE from *TBC1D32*-associated IRD (28). Hence, we assayed vascular endothelial growth factor (VEGF), which is primarily secreted basally, and pigment epithelium-derived factor (PEDF), which is primarily secreted apically (33), from control and *RLBP1* RPE. We did not detect significant differences in basal VEGF and apical PEDF secretion between control, RPA2 and RPA3 RPE (Fig 4H) but, interestingly, we detected a significantly elevated VEGF secretion apically in RPA3 RPE, and a significantly reduced PEDF secretion basally in RPA2 and RPA3 RPE, compared to controls.

Taken together, the RPE from *RLBP1* patients shows a clear retinosome accumulation, which could represent a potential therapeutic readout.

### AAV-RLBP1 encodes two CRALBP isoforms that are native to the mammalian RPE

We previously showed that AAV2/5 is the most effective serotype at transducing human RPE (26), thus we designed an AAV2/5 vector carrying *RLBP1* under control of the CAG promoter (AAV-RLBP1). To perform a proof-of-concept gene replacement study with clinical translatability, we subcloned the transgene cassette into two essentially identical proviral plasmids, pDDO and pKL; the latter being more suitable for clinical development due to the larger distance between the kanamycin resistance cassette and the 5’ inverted terminal repeat (ITR). Following transfection of the plasmids into COS-7 cells, we confirmed *RLBP1* expression by qPCR (Fig. 5A), and CRALBP expression by IF studies (Fig. 5B) and western blot analyses (Fig. 5C). Unexpectedly, we detected a specific signal migrating at the expected size (36 kDa) but composed of two bands (Fig. 5C), irrespective of the proviral plasmid and the intronless *RLBP1* cassette. To exclude a technical origin, we repeated western blot analysis under different lysis conditions (Fig. S2B) but in all cases two CRALBP bands were detected. Moreover, we tested denaturing versus native conditions using a mouse monoclonal or polyclonal anti-CRALBP antibody. Under denaturing conditions, two bands were detected with the monoclonal (Fig. S2C) and polyclonal (not shown) antibodies, although the latter gave high background. Under native conditions, the two CRALBP bands were highly resolved and more or less detectable, depending on the lysis buffer, using the polyclonal antibody (Fig. S2D) but not the monoclonal antibody (not shown).

**Figure 5:**
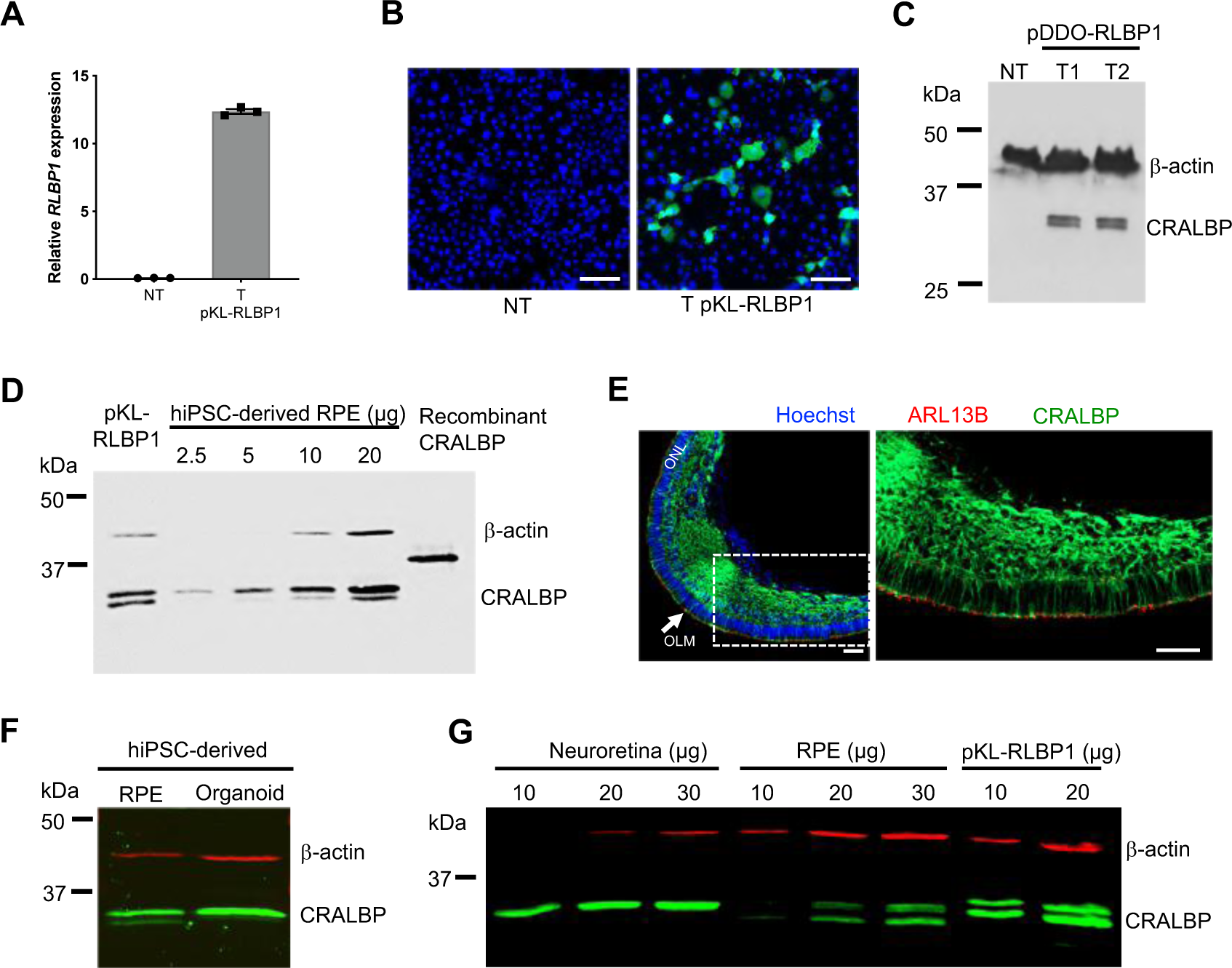
Unveiling of dual CRALBP isoforms. **A**) qPCR analysis of *RLBP1* expression, using a F primer spanning the exon 6 to 7 junction and a R primer in exon 7, in COS-7 non-transfected (NT) or transfected (T) with pKL-RLBP1. Data are represented as mean ± SEM. **B**) IF studies of CRALBP in NT and T COS-7 cells using a mouse monoclonal anti-CRALBP antibody. Scale bar = 50 µm. **C)** Western blot analysis of CRALBP using a mouse monoclonal antibody in NT and COS-7 cells transfected (T1 and T2) with pDDO-RLBP1 under standard conditions on an AnyKD gel. β-actin was the loading control. **D**) Western blot analysis of CRALBP in human iPSC-derived RPE at different loading levels on a 15% polyacrylamide gel. Positive controls, COS-7 cells transfected with pKL-RLBP1 and 200 ng recombinant CRALBP protein; β-actin was the loading control. **E**) CRALBP and ARL13B in human iPSC-derived retinal organoids at differentiation day 225; nuclei are labelled in blue; ONL, outer nuclear layer; OLM, outer limiting membrane (arrow). The right panel is a magnification of the region boxed in the left panel with only the red and green channels showing CRALBP expression in the Müller cells and ARL13B expression in the photoreceptors beyond the OLM formed by the Müller end feet. Scale bars = 50 µm. **F**) Western blot analysis of CRALBP in human iPSC-derived RPE and retinal organoids; β-actin was the loading control. **G**) Western blot analysis of CRALBP in the neuroretina and RPE of wild type mice; HEK-293 cells transfected with pKL-RLBP1 were used as a positive control for the double band profile; β-actin was used as the loading control.

To rule out that the two CRALBP isoforms were a consequence of *RLBP1* overexpression, we tested endogenous CRALBP by western blot analysis of healthy iPSC-derived RPE. We detected two bands of the same size as those observed in transfected cells, the intensity of which increased proportional to protein loading (Fig. 5D), confirming that the two isoforms were native to human RPE; the His-tagged recombinant CRALBP protein loaded as a control migrated at a higher molecular weight (predicted 37.4 kDa). We then asked if the two CRALBP isoforms were also present in human Müller cells of the neuroretina, which express *RLBP1*. We thus assayed healthy human iPSC-derived retinal organoids without RPE, in which CRALBP is localised in the prolongations of the Müller cells throughout the outer nuclear layer and in the Müller end feet forming the outer limiting membrane (Fig. 5E). We only detected the larger CRALBP isoform in the retinal organoids by western blot analysis, in contrast to the RPE (Fig. 5F). To further confirm that CRALBP expression in the human iPSC-derived retinal tissues was representative of the *in vivo* situation, we tested Cralbp expression in wildtype murine neuroretina and RPE separately. We clearly detected two Cralbp isoforms in the murine RPE and exclusively detected the larger isoform in the neuroretina (Fig. 5G), thus validating the results obtained in the human iPSC-derived models.

In conclusion, *RLBP1* expresses two CRALBP isoforms in the RPE, but only the larger canonical isoform in the neuroretina.

### Smaller CRALBP arises from an internal methionine and has a role in the visual cycle

To determine the origin of the smaller CRALBP isoform, we initiated mass spectrometry proteomic analysis of recombinant CRALBP and COS-7 cells transfected with pKL-RLBP1. Analysis of three gel fractions spanning the area of both isoforms detected various peptide fragments corresponding to CRALBP (Fig. S3A). Interestingly, N-terminal peptides containing the first nine aa of CRALBP were identified in only the 1^st^ gel fraction of highest MW, whereas N-terminal peptides beginning at a second methionine codon at aa position 10 were identified in the 2^nd^ and 3^rd^ fractions. These results suggested that differential initiation of translation from different methionine codons resulted in the two CRALBP isoforms. To confirm, we performed site-directed mutagenesis of the methionine to alanine codons in pKL-RLBP1 and generated pKL-RLBP1-Met1Ala mutated in the residue at position 1 and pKL-RLBP1-Met10Ala mutated in the residue at position 10. Following transfection in COS-7 cells and western blot analysis, we detected two CRALBP bands from the maternal plasmid (Fig. 6A) and exclusively detected the smaller band from pKL-RLBP1-Met1Ala, and the larger band from pKL-RLBP1-Met10Ala. Furthermore, IF studies of transfected HEK-293 cells detected CRALBP from each construct (Fig. 6B). In light of this data, we reanalysed the western blot analyses of the patient iPSC-derived RPE and noted that at higher loading levels with longer migration, two CRALBP bands were also detectable in the RPE of RPA2 (Fig. S3B); overexpression of the western blot of RPA1 RPE shown in Fig. 4G also suggested a second band (not shown). To unequivocally determine whether the mutant alleles c.25C>T and c.700C>T carried by RPA2 and RPA1, respectively, expressed both CRALBP isoforms, we introduced these variants into pKL-RLBP1 by site-directed mutagenesis, transfected HEK-293 cells and performed western blot analysis, which confirmed these observations (Fig. S3C).

**Figure 6:**
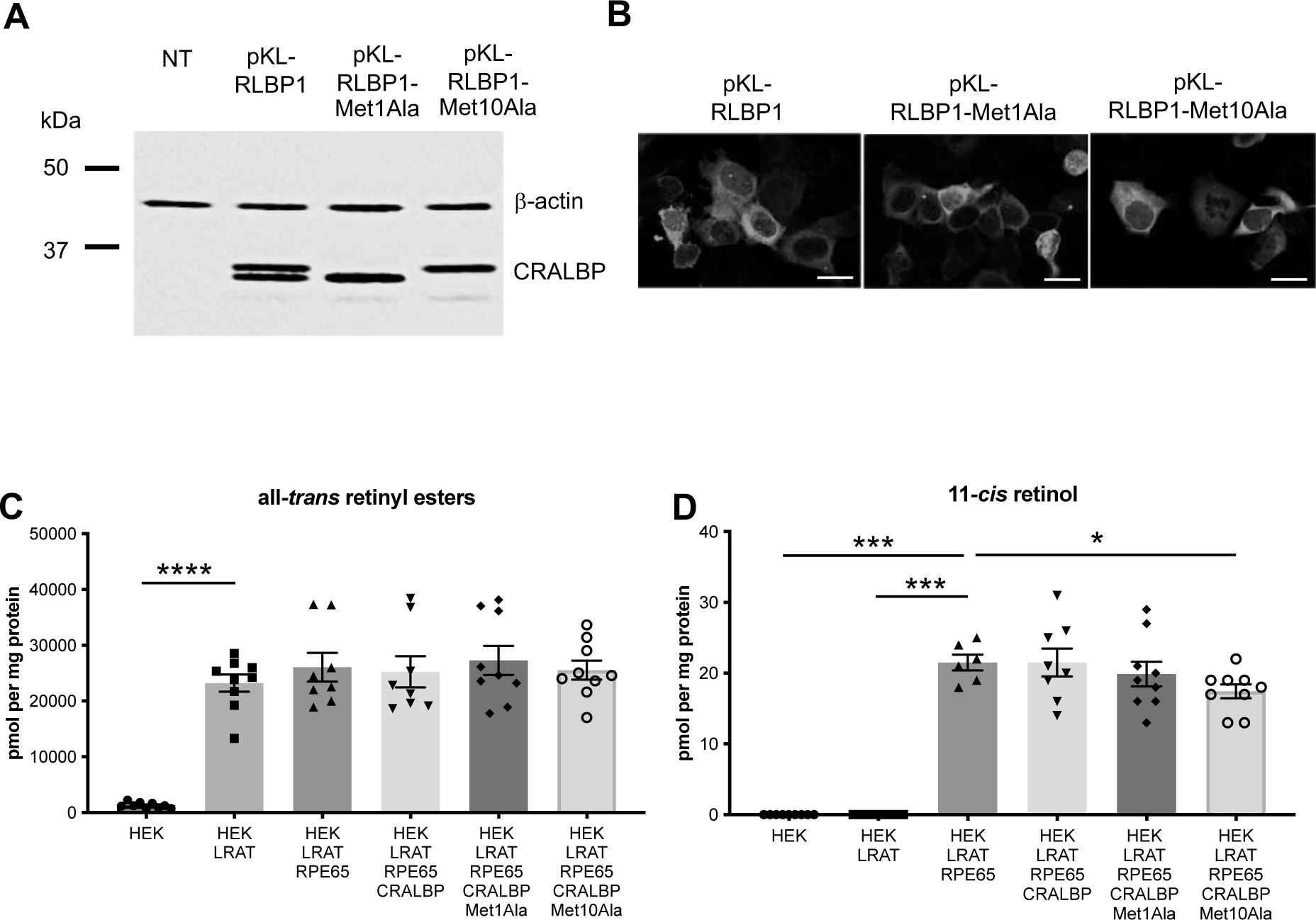
Origin and functional analysis of CRALBP isoforms. **A**) Western blot analysis of CRALBP in COS-7 cells non-transfected or transfected with the control or mutant (Met1Ala, Met10Ala) pKL-RLBP1 plasmids. Ten µg cell lysates were migrated on a 15% polyacrylamide gel; β-actin was used as the loading control. **B**) IF studies of CRALBP expression using a mouse monoclonal antibody in HEK-293 cells transfected with the control or mutant pKL-RLBP1. Scale bar = 20 µm. HPLC analysis of all-*trans* retinal esters (**C**) and11-*cis* retinol (**D**) levels (expressed in pmol per mg protein) in HEK-293 cells transfected with LRAT, RPE65 and/or control or mutant RLBP1 plasmids and incubated with all-*trans* retinol. Data are represented as mean ± SEM. \**p*<0.05; \*\*\**p*<0.001; \*\*\*\**p*<0.0001; *n* is indicated by the symbols.

In the visual cycle, CRALBP chaperones 11-*cis* retinol produced by the RPE65 isomerisation reaction to the RDH enzymes for oxidation thus enhancing RPE65 activity (34). To evaluate the impact of the two CRALBP isoforms on 11-*cis* retinol production, we transfected a stable LRAT-expressing HEK-293 cell line (35) with a plasmid expressing RPE65 alone or in combination with pKL-RLBP1, pKL-RLBP1-Met1Ala or pKL-RLBP1-Met10Ala. We then initiated an artificial visual cycle by adding all-*trans* retinol (vitamin A) to the transfected cells and assayed all-*trans* retinyl ester and 11-*cis* retinol production by HPLC. We confirmed that the LRAT-expressing cells produced significantly higher all-*trans* retinal esters levels compared to control cells (Fig. 6C) due to the esterification of the exogenous all-*trans* retinol. These levels were not significantly altered by the addition of the RPE65 or CRALBP plasmids. By comparison, 11-*cis* retinol levels were only detected upon addition of RPE65 due to the isomerisation reaction (Fig. 6D). They did not vary significantly in the presence of both CRALBP isoforms, but showed a tendency to decrease in the exclusive presence of the smaller isoform (Met1Ala) and were significantly reduced in the exclusive presence of the larger CRALBP isoform (Met10Ala).

Taken together, these results suggest that the smaller CRALBP isoform, which results from the differential use of a second methionine codon, also plays a role in the visual cycle.

### AAV-RLBP1 reduces retinosome accumulation in *RLBP1* iPSC-derived RPE

We next performed a proof-of-concept for AAV-RLBP in functionally validated, patient-specific iPSC-derived RPE and determined efficacy by assaying retinosome accumulation. We transduced the iPSC-derived RPE of patients RPA2 and RPA3 with AAV-RLBP1 and performed western blot analysis 1-mo post-transduction and clearly discerned two CRALBP bands (Fig. 7A). Furthermore, CRALBP levels significantly increased in RPA2 and RPA3 RPE post-transduction, compared to untreated tissue. Moreover, we assayed perilipin-2 levels by western blot analysis and detected significantly decreased levels in the RPA2 and RPA3 RPE post-AAV-RLBP1 transduction, compared to the untreated tissues (Fig. 7B). Lastly, we consolidated these results by performing a quantitative TEM study of the number of droplets in non-transduced and transduced RPA2 and RPA3 RPE and detected a significant decrease in the number of droplets following AAV-RLBP1 transduction (Fig. 7C).

**Figure 7:**
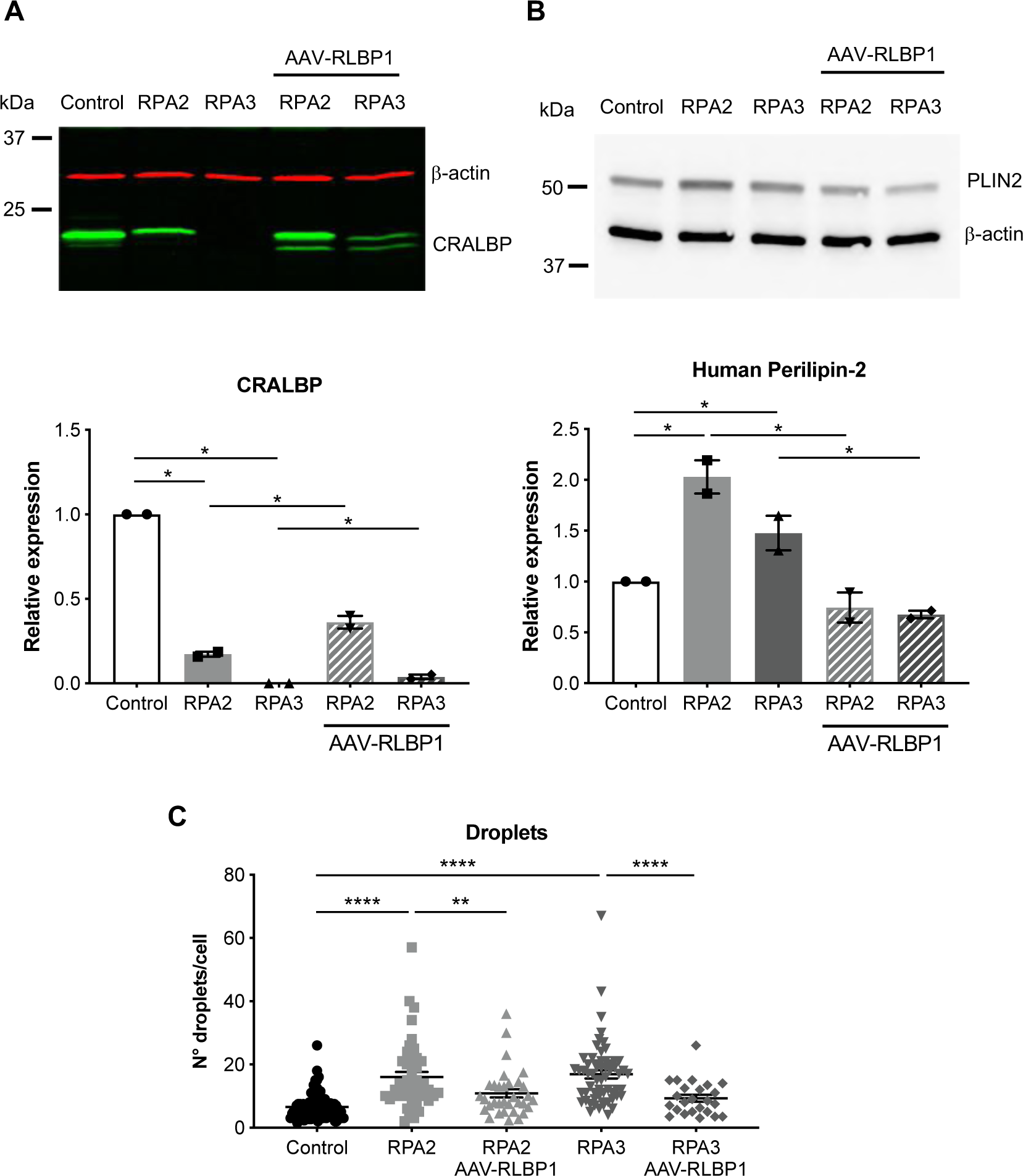
Rescue of retinosome accumulation in iPSC-derived RPE by AAV-RLBP1. **A**) Representative western blot analysis of CRALBP in control RPE and non-transduced and AAV-RLBP1-transduced RPA2 and RPA3 RPE; β-actin represents the loading control. The graph represents the quantification of CRALBP normalised to β-actin and expressed relative to control in 2 independent experiments. Data are represented as mean ± SEM; \**p*<0.05. **B**) Representative western blot analysis of PLIN2 in control RPE and non-transduced and AAV-RLBP1-transduced RPA2 and RPA3 RPE; β-actin represents the loading control. The graph represents the quantification of PLIN2 normalised to β-actin and expressed relative to control in 2 independent experiments. Data are represented as mean ± SEM; \**p*<0.05. **C**) Quantitative analysis of the number of droplets/cell following TEM analysis of control RPE and non-transduced and transduced RPA2 and RPA3 RPE. Data are represented as mean ± SEM; *p*<0.05; \*\**p*<0.01; \*\*\**p*<0.001; \*\*\*\**p*<0.0001; *n* is indicated by the symbols.

In conclusion, we provide the proof-of-concept of AAV-RLBP1 efficiency in a human context and demonstrate the pertinence of *RLBP1* iPSC-derived RPE for evaluating therapeutic efficiency.

### AAV-RLBP1 improves visual cycle kinetics and visual function in *Rlbp1^−/−^* mice

Lastly, we validated our *ex vivo* human AAV-RLBP1 proof-of-concept study on the *in vivo Rlbp1^−/−^* murine model, which was reported to show delayed visual cycle recovery following photobleaching accompanied by an accumulation of all-*trans* retinyl esters (16). To this end, we first crossed the albino *Rlbp1^−/−^* mice onto a pigmented strain to facilitate subretinal injections and verified *RLBP1* expression by qPCR (Fig. 8A) and CRALBP by western blot on low resolution AnykD gels (Fig. 8B) at 4-weeks post-transduction.

**Figure 8:**
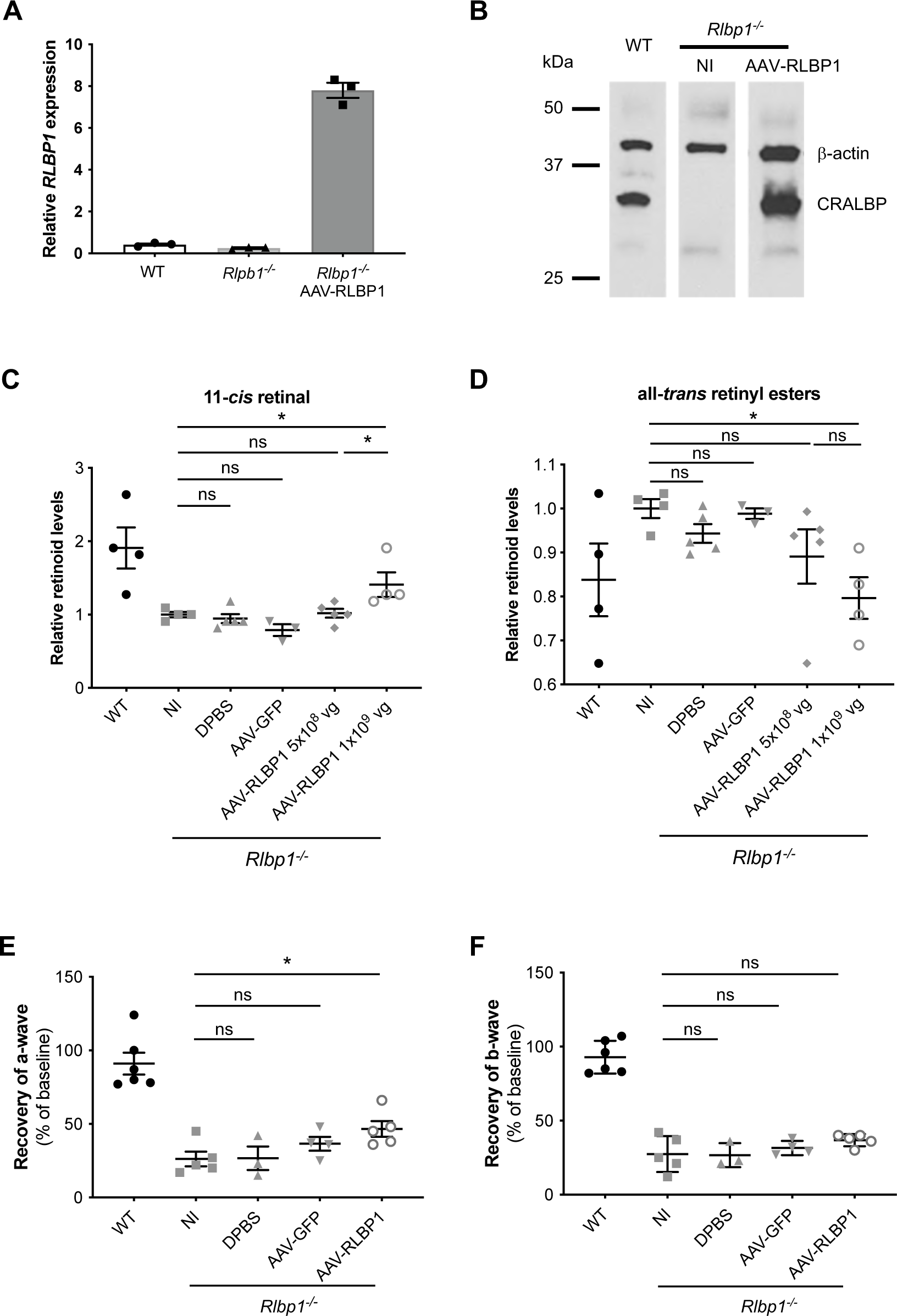
Rescue of visual cycle kinetics and function in *Rlbp1^−/−^* mice by AAV-RLBP1. **A)** qPCR analysis using the human-specific *RLBP1* exon 6 to 7 primers in the eyes of wild type (WT) and *Rlbp1^−/−^* mice non-injected (NI) or injected with 1×10^9^ vg of the AAV-RLBP1 vector at 4 weeks post-treatment. Data are represented as mean ± SEM. **B**) Representative western blot analysis of CRALBP in the eyes of WT and *Rlbp1^−/−^* mice NI or injected with AAV-RLBP1; β-actin represents the loading control. The mouse monoclonal anti-CRALBP antibody cross-reacted with murine CRALBP in the WT mouse. Under these migration conditions, the two CRALBP isoforms were not separated. HPLC analysis of 11-*cis* retinal (**C**) and all-*trans* retinal ester (**D**) levels in WT and *Rlbp1^−/−^*mice that were NI, injected with DPBS, AAV-GFP, or a low (5×10^8^ vg) or high (1×10^9^ vg) dose of AAV-RLBP1 at 8 weeks post-treatment. All data are represented as mean ± SEM and expressed relative to NI *Rlbp1^−/−^*mice; \**p*<0.05; ns, non-significant; *n* is indicated by the symbols. Analysis of the recovery of a-wave (**E**) and b-wave (**F**) ERG amplitudes (as a percentage of baseline amplitudes) at a light intensity stimulation of 1.5 log.cd.s.m^-2^ in WT and NI, DPBS-injected, AAV-GFP-injected and AAV-RLBP1 (1×10^9^ vg)-injected *Rlbp1^−/−^* mice at 10 weeks post-treatment. Data are represented as mean ± SEM; \**p*<0.05; ns, non-significant; *n* is indicated by the symbols.

We then assayed the recovery of visual cycle kinetics following AAV-RLBP1 treatment of *Rlbp1^−/−^* mice by HPLC analysis of all-*trans* retinyl ester and 11-*cis* retinal levels. Overall, mice were treated with three different doses, 5×10^8^ vg (Fig. 8), 1×10^9^ vg (Fig. 8 and Fig. S4) or 2×10^9^ vg (Fig. S4) of AAV-RLBP1; DPBS or equal doses of an AAV2/5-CAG vector expressing GFP (referred to as AAV-GFP) were administered as surgical and vector controls, respectively. Eight-weeks post-administration, all experimental groups were photobleached, dark-adapted, and the eyes assayed for retinoid content. Consistent with the albino strain (16), 11-*cis* retinal production levels were lower (Fig. 8C and Fig. S4A) and all-*trans* retinyl ester levels were higher (Fig. 8D and Fig. S4B) in pigmented NI *Rlbp1^−/−^* mice compared to wild type (WT). Following treatment with the intermediate 1×10^9^ vg (Fig. 8C and Fig. S4B) and high 2×10^9^vg (Fig. S4B) doses of AAV-RLBP1, we observed significantly increased 11-*cis* retinal levels compared to NI littermates, whereas levels were not significantly altered in DPBS- or AAV-GFP-injected controls (Fig. 8C). Similarly, we detected significantly decreased all-*trans* retinyl ester levels in *Rlbp1^−/−^* mice treated with 1×10^9^ vg (Fig. 8D and Fig. S4B) and 2×10^9^ vg (Fig. S4B) AAV-RLBP1 compared to NI littermates, whereas levels were not significantly altered in DPBS- or AAV-GFP-injected controls. By contrast, 11-*cis* retinal (Fig. 8C) and all-*trans* retinyl (Fig. 8D) levels were not significantly altered in mice treated with the lowest dose of 5×10^8^ vg AAV-RLBP1 compared to NI controls, demonstrating a dose-dependent effect. Lastly, as no significant differences in 11-*cis* retinal (Fig. S4A) or all-*trans* retinyl ester (Fig. S4B) levels were detected between the intermediate 1×10^9^ vg and high 2×10^9^ vg AAV-RLBP1 doses, we retained the 1×10^9^ vg dose for all subsequent experiments.

To determine that AAV-RLBP1-treated *Rlbp1^−/−^* mice also improved visual function, we assayed electroretinogram (ERG) recordings following light stimulation. To this end, 8-weeks post-subretinal injections, we first assayed baseline visual function of WT and NI, DPBS-injected, AAV-GFP-injected and AAV-RLBP1-injected *Rlbp1^−/−^*mice. As expected, all experimental groups showed similar a-wave (primarily generated by photoreceptors; Fig. S4C) and b-wave (primarily generated by bipolar cells; Fig. S4D) responses that decreased or increased, respectively with increasing light intensity. Then, 10-weeks post-treatment, we assayed the visual function by ERG recordings of the same animals following photobleaching and dark-adaptation. The AAV-RLBP1-injected mice showed a-wave (Fig. S4E) and b-wave (Fig. S4F) amplitude profiles that were closest to those of the WT mice, as compared to the NI and DPBS- or AAV-GFP-injected groups. To determine the recovery of visual function for each wave, the values post-photobleaching were expressed as a percentage of baseline values per mouse and averaged for each experimental group. WT mice almost fully recovered their a-wave (Fig. 8E) and b-wave (Fig. 8F) amplitudes following photobleaching and dark-adaptation whereas NI *Rlbp1^−/−^* mice did not. Similarly, there was no significant differences in the recovered a- or b-wave amplitudes in DPBS-injected or AAV-GFP-injected *Rlbp1^−/−^* mice compared to NI mice. By contrast, AAV-RLBP1-injected *Rlbp1^−/−^* mice showed significant improvement in a-wave amplitudes (Fig. 8E) compared to NI controls, but not in b-wave amplitudes (Fig. 8F).

Taken together, treatment by AAV-RLBP1 improves visual cycle kinetics and outer retinal function in *Rlbp1^−/−^* mice.

## Discussion

The advances made in the clinical translation of gene supplementation therapy for IRDs had provided much optimism for the patient and scientific community alike. Following the successful *RPE65* story, many clinical trials were initiated for diverse IRD genes (36), including *RLBP1* in 2018 (NCT03374657). We report here a novel proof-of-concept study for *RLBP1* supplementation therapy *ex vivo* in patient-specific iPSC-derived RPE models and *in vivo* in the murine knockout model. In addition to demonstrating phenotypic or functional restoration in both models, we discovered that *RLBP1* expresses two CRALBP isoforms. Moreover, we provide direct evidence that these isoforms are differentially expressed in both human and murine retina, and our data suggest that the small CRALBP isoform is necessary for optimal visual cycle activity.

Western blot analysis of CRALBP has been extensively described in various species (bovine (37), murine (38), human (39)) without suspicion of a second isoform. Some rare studies reported a second CRALBP band following western blotting analysis, but it was considered to be a product of degradation (40) or proteolysis (41). In one study, western blot analysis of HEK-293 cells transfected with *RLBP1*-expressing plasmids clearly resulted in two CRALBP bands, but these were not commented on by the authors (42). Similarly, western blot analyses of human ARPE19 cells overexpressing MITF (43) showed the presence of two CRALBP bands that were not highlighted. Particularly interesting was the study on human donor RPE that showed the presence of two CRALBP bands following western blot analysis of fresh RPE culture (44). Although uncommented by the authors, this study confirms our observations on human iPSC-derived RPE. In another encouraging example, Wenzel *et al*, analysed CRALBP, in parallel to RGR, RPE65 and RDH5, expression in murine eye cups, and the CRALBP bands were clearly thicker than those of the other proteins (45). Understandably, this observation was not highlighted by the authors but it clearly resembled our results on non-resolutive gels of murine RPE where the two CRALBP bands migrated close together.

Of our extensive literature search, only one study clearly speaks of two CRALBP bands following western blot analysis (46). The authors studied the role of cralbp in the visual cycle in zebrafish and attributed the two bands to the identification of two *rlbp1* ohnologues present in this species, in comparison to the “expected” single band in bovine and mouse eye. It should be mentioned though that the band corresponding to the bovine whole eye was quite thick, again raising the question as to whether it represented two bands in one. The zebrafish *rlbp1* ohnologues were referred to as *rlbp1a*, which is predominant in the RPE, and *rlbp1b*, which is expressed in Muller glia. The cralbp a isoform is slightly smaller and is missing the PDZ-domain target motif at the C-terminus that is present in cralbp b, suggesting distinct roles. By contrast, both human CRALBP isoforms contain the C-terminal PDZ-domain target motif but vary in the N-terminus. Along this line, Crabb *et al*, described the development of a recombinant human CRALBP protein and observed a doublet on SDS-PAGE analysis that was due to a minor truncation of the first 8 aa residues (47). It was concluded that the truncation of CRALBP might be due to lability of the N-terminal region to proteolysis. We did perform an *in silico* study of the CRALBP protein sequence using the N-terminal protein translational modification prediction program Terminus (http://terminus.unige.ch/). The only notable prediction was an N-alpha acetylation and cleavage of the first 5 amino acids (aa) of CRALBP (MSEGV). If the N-terminal region was cleaved or underwent proteolysis, this would also result in the differential use of the second N-terminal methionine initiation codon at aa 10, leading to the production of the 1-kDa smaller CRALBP isoform.

Using an *in vitro* visual cycle assay, we showed that 11-*cis* retinol levels tended to be reduced when one of the two CRALBP isoforms was absent and were significantly reduced when the small isoform was absent, suggesting a role in the visual cycle. CRALBP has been reported to have a dual role in the visual cycle by first chaperoning 11-*cis* retinol to the RDH enzymes for oxidation (48) and second chaperoning 11-*cis* retinal to the RPE membrane for binding to IRBP (41, 49). The way that CRALBP accomplishes these roles has not been fully elucidated. It is tempting to speculate that the two isoforms may have differential roles in the visual cycle with one isoform predominantly involved in retinol, and the other in retinal, binding. Interestingly, the small isoform is not needed in the Müller cells, as it was not detected in human or murine neuroretina. In the RPE visual cycle, CRALBP is required for the timely recovery of mammalian rod and cone responses following photobleaching (16). In the Müller visual cycle, CRALBP is needed to support cone-mediated vision and accelerate cone dark adaptation (21). Thus, it appears that there is no role for the smaller isoform in the cone-specific visual cycle. Given the reported cooperation between RGR and CRALBP in a photic visual cycle in both RPE and Müller glia (19), and the fact that 11-*cis* retinal is the sole product that would be produced therein further supports the rationale for the presence of only the longer isoform in Müller glia but both in RPE where 11-*cis* retinol is also present. A more detailed comparison of these two cycles could therefore yield clues to its role in the RPE.

As the two CRALBP isoforms appear to be essential in the RPE, it is therefore important, in terms of gene delivery, that the transgene cassette is capable of producing both isoforms. The two CRALBP isoforms were expressed from our AAV-RLBP1 vector and their presence likely contributed to the success of this proof-of-concept studies in both patient-specific iPSC-derived RPE and murine *Rlbp1^−/−^* RPE. It should be noted that the 11-*cis* retinal levels and a-wave recovery could not normalised to WT levels, as the bleb resulting from single administration subretinal injection only covered between 20-50% of the retina. By comparison, only a single CRALBP band was detected by western blot analysis following subretinal delivery of the previously reported scAAV2/8-hRLBP1-RLBP1 vector (25) that is currently being tested clinically. In order to rule out variables in sample preparation, we tested the same reported Cell Signalling buffer and protease inhibitors for the western blot analyses of our proviral plasmids, but we still detected two isoforms. It cannot be excluded that the single band following administration of the scAAV2/8-hRLBP1-RLBP1 vector was due to the migration conditions, as only a single CRALBP band was also expressed in the wild type mice. However, as the wild type CRALBP band was very faint, compared to the overexpressed exogenous CRALBP band, the second native CRALBP isoform may have been below the detection threshold.

An alternative explanation could lie with the different promoters used in the AAV2/5-CAG-RLBP1 and scAAV2/8-hRLBP1-RLBP1 vectors. Choi *et al* used a short endogenous *RLBP1* promoter that showed reduced *RLBP1* levels when compared to the long promoter in the single-strand vector genome conformation but reached similar levels in the self-complementary (sc) conformation (25). Thus, it could be hypothesised that the selected portion of the *RLBP1* promoter did not permit high enough transcript levels, in contrast to the strong ubiquitous CAG promoter used in our study, for both CRALBP isoforms to be detectable. If our conclusions concerning the importance of the second isoform are correct, then the efficiency of the scAAV2/8 vector in humans could be drawn into question and this will not be resolved until publication of the results of the corresponding clinical trial. Difficulties encountered with endogenous promoters was illustrated by another phase 1/2 clinical trial using an AAV2/2 vector with *RPE65* under control of its endogenous promoter (50) that did not have as dramatic results as the parallel study using a CAG promoter (51). This raised the hypothesis that the lower response could have been due to the use of a weaker promoter (52). Furthermore, Kennedy *et al.* have suggested that alternate promoter/elements might be used for CRALBP expression in Müller glia as opposed to RPE, but these have not been identified to date (53). It is thus conceivable that minimal regions of endogenous promoters may lack specific elements that could impact transcriptional regulation and the production of different protein isoforms, thus raising important considerations for gene supplementation therapy.

Another important aspect of our study is that, to our knowledge, we describe the first human iPSC-derived RPE models of *RLBP1*-associated IRDs and demonstrate salient genotype-phenotype correlations. We show that the iPSC-derived RPE from *RLBP1* patients expresses characteristic markers and remains tight in culture, similar to control RPE. The visual cycle defect emblematic of CRALBP deficiency was evidenced by increased all-*trans* retinyl ester levels and retinosome accumulation, which correlated with clinical severity. Similarly, we identified altered RPE functionality that was more evident in the RPE from patient RPA3 who presented with the severe NFRCD. As MERTK expression seemed intact, we speculate that the altered phagocytic activity, as well as growth factor secretion, were not primary defects but likely consequences of the subcellular changes due to retinosome accumulation. CRALBP expression in iPSC-derived RPE also varied depending on the *RLBP1* variant/clinical form. In control iPSC-derived RPE, CRALBP shows a dual subcellular localisation, and it is tempting to speculate that the cytosolic localisation could be related to its role in binding 11-*cis* retinol following photoisomerization, and the membrane localisation could be linked to the transfer of the 11-*cis* retinal to IRBP at the plasma membrane. This is consistent with prior localisation studies in mouse and rat RPE (54). Interestingly, in the RPE of patient RPA1 (BD), the plasma membrane localisation was lost and CRALBP appeared more cytosolic, which was similar to our previous observations in an iPSC-derived RPE model of another IRD due to TBC1D32 deficiency that also showed a disruption of the visual cycle (28). By contrast, in the RPE of patient RPA2 (RPA), presenting with the mildest clinical form, the dual CRALBP localisation pattern was preserved but CRALBP staining was less intense; no CRALBP expression, irrespective of isoform, was detected in RPA3, consistent with the severe NFRCD phenotype.

Both RPA1 and RPA2 RPE harbour a nonsense variant, p.Tyr111*, in combination with a missense variant, which we previously modelled (6). The reduced *RLBP1* levels in the RPE of these patients strongly suggest that the p.Tyr111* variant does not result in CRALBP expression. In RPA2 RPE, there was a clear expression of two CRALBP isoforms by western blot analysis. Therefore, we hypothesise that the pArg9Cys variant carried by RPA2, that is located immediately upstream of the second initiation codon, may allow production of a mutant larger isoform but an intact smaller isoform and thus account for the milder phenotype in this patient. This is supported by our *in vitro* visual cycle assay showing that the presence of the smaller isoform alone does not result in significantly decreased 11-*cis* retinol levels. By contrast, in RPA1 RPE, both CRALBP isoforms would carry the p.Arg234Trp mutant consistent with a more severe clinical phenotype. The crystal structure of CRALBP-p.Arg234Trp suggests that the mutant protein(s) has tighter binding of 11-*cis* retinal (55), which results in an increased resistance to light-induced photoisomerization (56). Interestingly, we less readily detected the smaller CRALBP isoform by western blot analysis in RPA1 RPE and, if we correlate this observation with the IF data showing the loss of CRALBP near the plasma membrane, it could suggest that the smaller isoform is associated with the membrane and hence, as speculated above, is involved in 11-*cis* retinal binding and transfer to IRBP for entry to the POS.

Taken together, further study of both control and patient-specific iPSC-derived RPE may yield insights into CRALBP function and the role the canonical and non-canonical isoforms in the visual cycle. In the meantime, the modelling of *RLBP1*-associated IRDs using iPSC-derived models allowed us to identify quantitative criteria to evaluate the efficacity of *RLBP1* gene replacement in human RPE, which was consolidated by our murine proof-of-concept study. Thus, these models will also be invaluable for evaluating other emerging therapies for *RLBP1*-associated IRDs.

## MATERIAL AND METHODS

### Clinical examinations and iPSC generation

Color fundus photography was performed using an automated, non-mydriatic fundus camera (AFC 330, Nidek). Dermal fibroblasts were cultured from skin biopsies and reprogrammed under feeder-free conditions using the integration-free CytoTune iPSC 2.0 Sendai Reprogramming kit (29). iPSCs were cultured on dishes coated with 1:100 dilution of Matrigel hESC-qualified Matrix (Corning) in Essential 8 (E8) medium (Gibco) and passaged weekly using 0.48 mM Versene solution (Gibco). For karyotype analyses, iPSCs were prepared as described (29) and twenty metaphase spreads were analysed using standard G-banding techniques (Chromostem facility, Montpellier University Hospital, France). For teratoma analyses, a confluent non-differentiated 35 cm^2^ dish of iPSCs was dissociated with Versene, centrifuged 5 min at 800 rpm, resuspended in E8 medium containing 30% Matrigel and injected into NOD.Cg-Prkdc^scid^/J mice (26). Paraffin-embedded 4-µm sections were stained with Hematoxylin-Eosin-Saffron using standard protocols and images taken using an upright Eclipse Ci-L plus microscope connected to a DS-Fi3 digital microscope camera and NIS-Elements L Imaging software (Nikon).

### iPSC-derived retinal differentiation

iPSCs were spontaneously differentiated into RPE and cultured in KnockOut DMEM medium (Gibco) supplemented with 20% KnockOut Serum Replacement (KOSR; Gibco), 1% GlutaMax (Gibco), 1% non-essential amino acids (Gibco) 0.1% β-mercaptoethanol and 1% penicillin-streptomycin (29). At passage (P) 3, RPE was seeded at a density of 6×10^4^ cells per 0.32 cm^2^ on a 1:30 dilution of Corning Matrigel hESC-qualified matrix on cell culture inserts with high density 0.4 µm pores (Falcon), 24-well plates or 6-well plates depending on the experiment. Trans-epithelial resistance (TER) measurements were performed as described without modification (29). iPSCs were differentiated into retinal organoids using a 2D-3D differentiation protocol supplemented with retinoic acid (57).

### Phagocytosis and growth factor secretion

Bovine photoreceptor outer segments (POS) were prepared as described (58). iPSC-derived RPE cultured on membrane inserts were fed with 7.5 POS/cell for 2.5 h at 37 °C, washed, dissociated with 0.25% trypsin and pelleted. Cells were resuspended in 200 µl PBS and fluorescence analysed on a BD Accuri C6 Flow Cytomoter (BD Biosciences). VEGF and PEDF levels were assayed by ELISA (R&D systems) on 24-hour conditioned media collected from the apical and basolateral chambers of 4 month-old iPSC-derived RPE cultured on inserts (33). Optical densities were determined at 450 nm using a microtiter plate reader (Clariostar; BMG Labtech). Samples were collected from 9 inserts per condition, assayed in triplicate and the mean expressed as percentage of control values.

### TEM and multiphoton microscopy

Three to 7 month-old iPSC-derived RPE cultured on inserts was processed and embedded for TEM analysis as described (29). Counterstained 70-nm sections were observed using a Tecnai F20 transmission electron microscope at 200 kV (CoMET facility, INM). Lipid droplets were counted manually and counting confirmed by an additional masked investigator. Fixed 7-month-old iPSC-derived RPE cultured on inserts was imaged using a Zeiss LSM 880 multiphoton associated with an Upright AxioExaminer microscope and equipped with a Ti:sapphire femtosecond laser (Chameleon Ultra II; Coherent France). Stimulation was performed at a wavelength of 740 nm and emission was recorded between 500 and 550 nm. Images were treated using the Imaris IsoSurface module (Oxford Instruments). The area of fluorescence was calculated relative to the area of each image.

### AAV proviral plasmid and vector generation

The *RLBP1* cDNA (NM_000326) from the ATG (+1) to the TGA codon (kindly provided by John Crabb, Cleveland Clinic Foundation) was PCR amplified using AmpliTaq Gold DNA polymerase (Applied biosystems) with the forward (F) *Mlu*I-containing primer and the reverse (R) *Xho*I-containing primer (see Table S1) under standard conditions. The 966-bp amplicon was purified using the Wizard SV Gel and PCR Clean-Up System (Promega) and subcloned into the TA cloning vector pGEM-T easy (Promega), according to the manufacturers’ recommendations. DNA was isolated from *RLBP1*-containing pGEM-T easy colonies and digested with *Mlu*I and *Xho*I (Promega) for 3 h at 37 °C. The excised *RLBP1* insert was gel purified using the Mini Elute Gel Extraction kit (Qiagen) and ligated using T4 DNA ligase (Promega) at RT for 1 h into the *Mlu*I/*Xho*I-digested recombinant AAV proviral plasmids pDDO-020-SSV9-CAG-bGHpA or pKL-AAV-CAG-bGHpA (TaRGet vector core, Nantes, France). Five µl of the ligation reaction were transformed into MAX Efficiency Stbl2 Competent Cells (Invitrogen) and cultured at 30 °C to minimise recombination between the ITRs. Resulting clones were screened by restriction enzyme digestion to assay for ITR stability and correct *RLBP1* insertion. Positive pDDO-RLBP1 or pKL-RLBP1 clones were verified by Sanger sequencing with a F primer in exon 5 and a R primer spanning the junction of exons 7 and 8. DNA from the selected clone was purified using the EndoFree Plasmid kit (Promega) and sent to the TaRGet vector core to generate the corresponding AAV-RLBP1 vector (production titres 1.2×10^12^ vg/ml (pDDO batch in DPBS 1X without CaCl_2_ and MgCl_2_; Gibco) and 3.3×10^12^ vg/ml (pKL batch in clinically-compatible ocular buffer).

### PCR analysis and Sanger sequencing

Plasmid DNA was isolated using the Qiaprep Spin Miniprep Kit (Qiagen) and genomic DNA using the DNeasy Blood and Tissue Kit (Qiagen). The exons containing the *RLBP1* variants carried by the patients RPA1 and RPA2 in exons 4, 5 and 8 were PCR amplified using the primers listed in Table S1. Amplicons were cleaned with the ExoSAP-IT PCR clean-up kit (GE Healthcare) and Sanger sequencing was performed using the BigDye Terminator Cycle Sequencing Ready Reaction kit V3.1 on a 3130xL Genetic Analyser (Applied Biosystems, Foster City, CA). A total of 50 ng amplified genomic DNA or 250 ng plasmid DNA and 3.2 µM of primer were used for each reaction. Reactions were precipitated and resuspended in 15 µl ultrapure formamide before sequencing. To assay for the large deletion carried by patient RPA3, long-range PCR was performed using the TaKaRa Taq DNA polymerase and a F primer in intron 6 of *RLBP1* and a R primer situated in the intergenic region between *RLBP1* and *ABHD2* gene (Table S1; (3)). Amplicons were migrated on a 1% agarose gel.

### Transfections

COS-7 and HEK-293 cells were plated at a density of 4×10^4^ cells/cm^2^ or 1×10^5^ cells/cm^2^, respectively, in 24-well plates for qPCR or western blot analyses, on glass coverslips in 24-well plates for IF studies, in 12-well plates for western blot analysis or in 6-well plates for retinoid assays. Cells were cultured in DMEM media supplemented with 10% foetal bovine serum (Gibco) for 24 h prior to transfection. Experimental wells were transfected with 500 ng (24-well plates), 1 µg (12-well plates) or 2.5 µg (6-well plates) of plasmid DNA diluted in OptiMEM medium using the Lipofectamine 3000 reagent (Invitrogen, ThermoFisher Scientific, France) for 48 h. As a control condition, non-transfected cells were incubated with the transfection reagents in Opti-MEM medium without plasmid DNA. For all experiments, transfections were performed in duplicate.

### Proteomic analysis

Transfected COS-7 cells were scraped in ice cold PBS containing Complete protein inhibitor cocktail tablets (Roche), centrifuged at 200 x g for 5 minutes, and the pellet was then lysed in RIPA buffer (5 M NaCl, 0.5 M EDTA pH 8, 1 M Tris pH 8, 10% sodium deoxycholate, 10% SDS) under agitation at 4 °C for 30 min. The lysate was cleared by centrifugation at 21 130 x g at 4 °C for 30 min and protein content quantified using the Pierce BCA protein assay kit (ThermoFisher Scientfic). Twenty µg of the lysate, 2.5 µg of a recombinant His-tagged CRALBP protein (ab177594, Abcam) and a SeeBlue Prestained Protein marker (LC59225, ThermoFisher Scientific) were loaded on two 12% precast gels (Life Technologies system). Following migration, one gel was transferred onto a PVDF membrane and hybridised with the mouse monoclonal anti-CRALBP antibody. The second gel was rinsed and stained in Coomassie blue staining solution (InstantBlue, Expedeon) for 1 h. The bands of interest were cut from the Coomassie stained gel using the western blot filter as a guide and an in-gel digestion using a protease (Sequencing grade Trypsin, Promega) was performed prior to proteomic analysis by nanoscale liquid chromatography coupled to tandem mass spectrometry (nano LC-MS/MS) using a QTOF Impact II (Bruker Daltonics) mass spectrometer, as described (59).

### Site-directed mutagenesis

The ATG (+1) codon of pKL-RLBP1 was mutated to an alanine codon (GCG) using the QuikChange Site-directed Mutagenesis kit (Agilent technologies) with the F primer and its reverse complement (Table S1) to generate pKL-RLBP1-Met1Ala. The ATG (+10) codon of pKL-RLBP1 was mutated to a GCG codon using the Assembly PCR oligo maker (60). A first round of amplification was performed for 8 cycles using a mix of six overlapping primers (Table S1) spanning the first five *RLBP1* exons (primer F1 flanked the ATG (+1) codon and R2 contained the mutated ATG). A second round of amplification was performed for 25 cycles using primers flanking the ATG codon (22 bp upstream and 200 bp downsteam). The purified amplicon was digested by *Mlu*I and *Bsa*I and subcloned into *Mlu*I/*Bsa*I-digested pKL-RLBP1 to generate pKL-RLBP1-Met10Ala. Following mutagenesis and/or subcloning, both ATG-mutated plasmids were transformed into MAX Efficiency Stbl2 Competent Cells (Invitrogen). Clones were screened by restriction enzyme digestion and verified by Sanger sequencing using F primers in the pKL backbone upstream of *RLBP1* and in exons 5 and 7 of *RLBP1,* and R primers in exons 5 and 6 of *RLBP1* (Table S1), to cover the entire open reading frame. Prior to the generation of the c.700C>T (p.R234W) and c.25C>T (p.R9C) mutagenesis plasmids, the *RLBP1* cDNA was removed by *Mlu*I/*Xho*I digestion of pKL-RLBP1, and subcloned into pGEM-T easy to avoid unwanted recombination events due to the repetitive ITR sequences. Mutagenesis was performed using the primers listed in Table S1 and verified by Sanger sequencing. The mutant cDNAs were subcloned back into the pKL plasmid to generate pKL-RLBP1-Arg9Cys and -Arg234Trp.

### Reverse Transcription (RT) and Quantitative (q) PCR analyses

Transfected COS-7 cells and iPSC-derived RPE were dissociated with 0.25% trypsin (Gibco), and iPSCs with Versene, centrifuged and snap-frozen in liquid nitrogen. Following sacrifice, WT and untreated and treated *Rlbp1^−/−^*mouse eyes were enucleated, the anterior segment and the lens were removed, the neuroretina dissected, and the remaining RPE scraped and collected. The neuroretina and RPE were pooled and snap-frozen. RNA from transfected cells, iPSCs, iPSC-derived RPE, and mouse tissues were isolated using the QiaShredder and RNeasy mini kits (Qiagen, France). The isolated RNA was treated with RNase-Free DNase (Qiagen) and 150 ng or 500 ng (for iPSCs) was reverse transcribed using the Superscript III Reverse Transcriptase kit (Life Technologies, ThermoFisher Scientific). Clearance of the Sendai vectors was assayed by RT-PCR amplification of fibroblast and iPSC cDNA with primers specific to the vector backbone (SEV), polycistronic cassette (KOS) and monocistronic cassettes (KLF4, c-MYC) (Table S2). qPCR amplification of a 1:10 dilution of cDNA was performed in triplicate on a LightCycler^®^ 480 II thermal cycler (Roche, France) using the LightCycler^®^ 480 SYBR Green I Master mix and specific primers (Table S2). Results were analysed using LightCycler^®^ 480 software and the Microsoft Excel programme Quantification was performed using the ΔΔCt method, normalised to *GAPDH* for human cells and *L27* for murine tissues, and expressed relative to control.

### Western blot analysis

Non-transfected and transfected COS-7 and HEK-293 cells, non-transduced and transduced iPSC-derived RPE, and organoids were scraped or collected in ice-cold PBS containing Complete protein inhibitor cocktail tablets (Roche) (unless otherwise stated) and centrifuged at 200 g for 5 minutes. Pellets were resuspended in 30 µL 2x Laemmli sample buffer (Biorad, France) containing 1:25 dilution of ß-mercaptoethanol (Sigma-Aldrich) and 1 µl Benzonase (Sigma-Aldrich) or in RIPA buffer (unless otherwise stated) and centrifuged at 20 000 g for 15 min prior to protein quantification using the Pierce BCA protein assay. iPSC-derived organoids and dissected neuroretina and RPE from WT and *Rlbp1^−/−^* mice were lysed in 30 µl RIPA buffer using a mortar and pestle, centrifuged at 20 000 g for 15 minutes, and the proteins were quantified prior to loading (murine tissues) or directly loaded (organoids). Samples (25 µl total volume) were heated 5 minutes at 95 °C and loaded onto an AnyKD precast MiniProtean TGX Stain Free gel (Biorad) or an 15% SDS-PAGE gel, as specified. The separated proteins were electrotransferred using a Trans-Blot^®^ Turbo™ Mini PVDF Transfer Pack and System (Biorad). After blocking for 1 h in 0.5% Tween-PBS in 5% skim milk (blocking solution), membranes were incubated with the primary antibodies overnight at 4 °C, washed in 0.5%Tween-PBS, and incubated with the secondary antibodies 45 min at room temperature (RT). The detection step was performed using the Amersham ECL prime western blotting detection reagent (GE Healthcare, France) and visualised using autoradiographic film exposure and a Biorad ChemiDoc XRS+ Imager system or using an Odyssey CLx Imager (Li-COR Biosciences) and quantified with the Image Studio Lite or Empiria Studio software.

### Immunofluorescence studies

Uncoated or poly-D-lysine-coated (HEK-293) glass coverslips containing non-transfected and transfected COS-7 and HEK-293 cells, respectively, and inserts containing iPSC-derived RPE were fixed in 4% paraformaldehyde (PFA; Alfa Aesar) for 10 min at RT, blocked in 10% donkey serum (Millipore) and 1% BSA (Sigma-Aldrich) in PBS, and permeabilised with 0.3% Triton X-100 (Sigma-Aldrich). Retinal organoids were washed in PBS, fixed in 4% PFA, incubated in 30% sucrose/PBS overnight at 4 °C and embedded in Tissue-Tek OCT compound (Sakura). Slides with 10 µm cryosections were incubated in 10% donkey serum, 5% BSA and 0.1%Triton X100 for 1 h at RT. For all cell types, primary antibodies were incubated overnight at 4 °C and secondary antibodies with 0.2 mg/ml bisBenzimide Hoechst (Sigma-Aldrich) or 1 µg/ml DAPI (Sigma-Aldrich) were incubated 45 min at RT prior to mounting in Dako Fluorescent Mounting Media (Dako France SAS, Les Ulis, France). Cells were visualised using an ApoTome 2 Upright wide-field or a Confocal LSM 880 microscope (Carl Zeiss SAS).

### Visual cycle experiments

HEK-293 cells stably expressing human LRAT (35) were transfected with plasmids carrying control *RPE65* (35) and *RLBP1* sequences, or the mutated Met1Ala and Met10Ala *RLBP1* sequences. Under dim light, transfected HEK-293 cells were starved in serum-free medium for 8 h, and incubated for 24 h in KnockOut DMEM, 15% KOSR, 2% fatty acid-free BSA (Sigma) and 10 µM Vitamin A (Sigma, R7632). Human iPSC-derived RPE aged 6 to 8 weeks was starved in RPE medium without KOSR for 8 h and incubated for 24 h. After 8 h, the media was changed to RPE media without KOSR supplemented with 2% fatty acid-free BSA, 15% FBS (Gibco) and 10 µM Vitamin A (61). All cells were scraped in PBS, pelleted and stored in the dark at −80 °C. HEK-293 and RPE pellets were lysed in 200 µl of 0.2% SDS/PBS. Ten µl was withdrawn for protein quantification using the Pierce BCA protein assay (ThermoFisher) and 300 ml ethanol and 1 ml of hexane were added to the remainder of the lysate, which was then processed for HPLC analysis, as described below. Three independent experiments were performed.

### AAV transduction

Ten to 12 weeks post-seeding, iPSC-derived RPE was transduced with 50 000 vg/cell of AAV-RLBP1 in 100 µl (inserts) or 200 µl (24-well plates) of media for 6 h. The wells were then supplemented with 100 µl (inserts) or 200 µl (24-well plates) media overnight and another 100 µl or 500 µl, respectively, the next morning. The media was refreshed 2 days post-transduction and the transduced RPE was kept in culture for 8 weeks.

### *Rlbp1^−/−^* mouse colony

The *Rlbp1^−/−^* mouse model (16) of mixed 129/SvJ, C57BL/6J and BALB/c backgrounds was kindly provided by the Cleveland Clinic (Beachwood, OH, USA). The founders were crossed with C57BL/6J mice (Harlan France SARL, Gannat, France) and the colony maintained in a controlled environment with a 12-/12-hour light/dark cycle according to the European guidelines for the care and use of laboratory animals (EU directive 2010/63/EU). For genotyping, genomic DNA was extracted and the wild type and *Rlbp1^−/−^* alleles amplified using the Phire Tissue Direct PCR master mix (ThermoFisher Scientific) and a forward primer in combination with 2 reverse primers (Table S1) to separate the WT and *Rlbp1^−/−^*alleles (16).

### Subretinal injection

Mice were anesthetised with ketamine (70 mg/kg; Merial, France) and xylazine (28 mg/kg; Bayer Healthcare, France) systemically and oxybuprocaine 0.4% (Cebesine, Bausch + Lomb, France) locally. The pupils were dilated by the sequential administration of a drop of phenylephrine 10% (Neosynephrine, Europhta) and tropicamide 0.5% (Mydriaticum, Théa, France), and the cornea covered with a drop of Lacryvisc (Alcon, France) and a glass coverslip. Under a surgical microscope, the eye was first pierced at the corneal-scleral junction to relieve the intra-ocular pressure. Subsequently, subretinal injections were performed using a 5 µL Hamilton syringe and a bevelled 34 G needle. On the experimental day, the AAV-RLBP1 vector was diluted in DPBS or ocular buffer to the required dose and 2 µL was injected into one eye of a *Rlbp1^−/−^* mouse. The contralateral eye was either left non-injected or injected with 2 µl DPBS or an AAV-GFP vector to control for the surgical procedure and non-specific effects of the AAV vector, respectively. At the required time post-injection, animals were euthanised by cervical dislocation.

### High Performance Liquid Chromatography

Eight-weeks post-injection, the experimental mouse groups were photobleached at 6000 lux for 20 min and then dark-adapted for 4 h. The animals were then euthanised and the ocular globe rapidly enucleated. The eyes were homogenised in 300 µl of 3 M formaldehyde (Sigma), incubated at 30 °C for 5 minutes, and supplemented with 0.3 ml of ethanol (62). One ml of hexane (Sigma) was then added, the samples homogenised and the organic phase collected. This step was repeated. The samples were then concentrated to 20 µl using a Speed Vac vacuum concentrator. Analyses were performed using a Varian HPLC system equipped with a NUCLEODUR SiOH column (4.6 mm x250 mm) (Macherey-Nagel) and a Prostar 330 diode array detector. The elution was performed with 6% ethyl acetate in hexane for 10 min at a flow rate of 2ml/min. The retinoids were quantified from the peak areas using calibration curves determined with established standards.

### Electroretinogram recordings

All ERG recordings were performed following overnight dark adaptation, at the same time of the day, under dim red light in a dark room, and using the Visiosystem (SIEM, France) with cotton wick electrodes (63). Each animal was anesthetized, its pupils dilated, and kept on a heating pad throughout the experiment maintaining the rectal temperature at 37 °C (Temperature Control Unit HB 101/2; Bioseb, France). A ground needle electrode was placed subcutaneously near the tail, and 2 reference electrodes were placed near the ears. ERG responses were recorded simultaneously from both eyes. The response to seven flashes were averaged for each light intensity (−1, −0.7, −0.3, 0, 0.7, 1, 1.5 and 2 log candela second/metre^2^ (cd.s.m-^2^)). The duration of each flash was 5 ms with a frequency of 0.3 Hz. Baseline visual function of the mice in all experimental groups was measured 8-weeks post-treatment. At 10-weeks post-treatment, the same mice were dark-adapted overnight, their pupils dilated, and photobleached by exposure to 2000 lux for 5 minutes. Following illumination, mice were again dark-adapted for 4 h prior to ERG recordings. The percentage of recovery was determined as the ERG recording post-illumination over the baseline recording per eye. The values at a flash intensity of 1.5 cd.s.m^-2^ were used to generate the dot plots as the recordings were high but had not plateaued.

### Statistical analyses

Statistical analyses were performed using a 2-tailed Mann and Whitney test with GraphPad Prism 8.4 software (GraphPad Software, La Jolla California USA). A *p* value <0.05 was considered significant. The number of samples per experiment and condition are indicated in the corresponding figures or legends.

### Study approval

Clinical and genetic investigations were performed following signed informed consent in accordance with approved protocols of the Montpellier University Hospital (ID n°: IRB-MTP_2021_11_202100959) and in agreement with the Declaration of Helsinki. Skin biopsies and iPSC reprogramming were performed following ethics approval provided by the French National Agency for the Safety of Medicines and Health Products (ID n°: 2014-A00549-38). Animal care and experimentation were approved by local and national ethics committees under the project number APAFIS#1192-2015071713533208.

## Data availability

All data are available in the main text or the supplementary materials.

## Author contributions

Co-first authors were assigned by order of arrival on the project. KD, GD, LG, DM, MP, NE, CSS, FB, JV performed experiments. BB and IM provided genetic and clinical data. HB and CH analysed data. MR and YA shared expertise. PB and VK provided supervision and analysed results. VK provided funding and wrote the manuscript. All authors reviewed and edited the manuscript.

## Acknowledgements

We dedicated this work to Pr. Christian Hamel who helped initiate the study and passed away prematurely in 2017. We thank A. Conscience, M. Cavalier, A. Jemi, Z. Jazouli, K. Bellouma, C. Martin and T. Duncan for technical assistance, and A.F. Roux for critical reading of the manuscript. We also thank C. Cazevieille of the CoMET facility (INM), P. Clair of the for qPHD facility (University of Montpellier), the MRI imaging facility and the RAM-Neuro animal facility. We particularly thank S. Marconi and N. Delauney (formerly of Horama) for stimulating scientific discussions. We are grateful to the individuals who participated in the study. This work was funded by Retina France, AFM, Fondation de France and Fondation pour la Recherche Medicale. Part of the work was funded as a maturation project by the axLR SATT and a collaboration contract with Horama (now Coave Therapeutics).

