## Supplemental Figures and Tables for "Dual CRALBP isoforms unveiled: iPSC-derived retinal modelling and AAV2/5-RLBP1 gene transfer raise considerations for effective therapy"

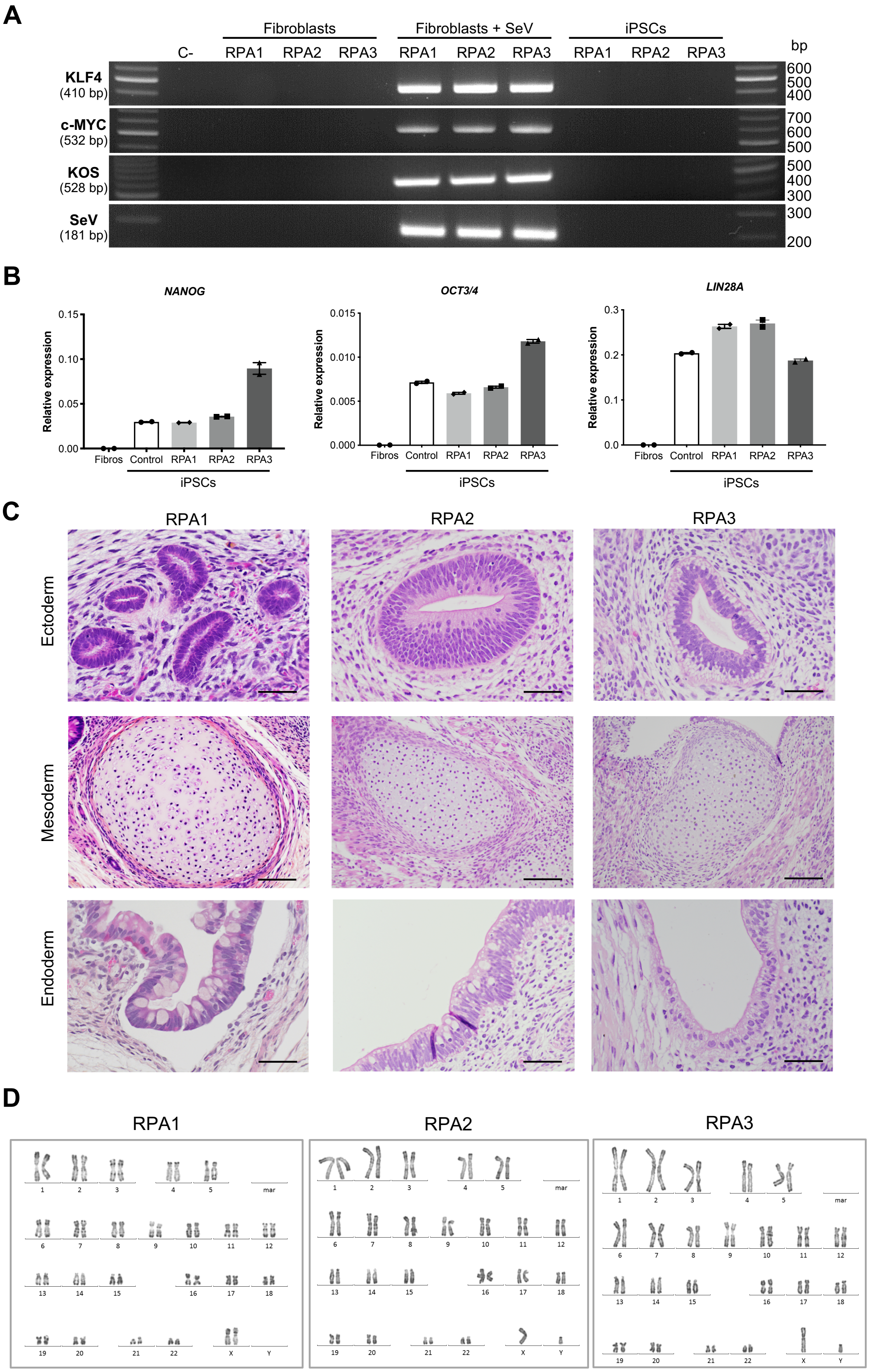


**Figure S1: Quality controls of RPA1, RPA2 and RPA3 iPSCs. A**) RT-PCR analysis of the clearance of the Sendai virus (SeV) reprogramming vectors in iPSCs at P12 (RPA2) and P16 (RPA1 and RPA3) using primers specific to the transgene cassettes (KLF4, c-MYC, KOS) or viral backbone (SeV). C-, control without cDNA; Fibroblasts, negative control; SeV-transduced fibroblasts (+ SeV), positive control. **B**) qPCR analysis of the relative expression of host pluripotency genes *NANOG*, *OCT3/4* and *LIN28A* in control iPSCs (positive control) and iPSCs of RPA1, RPA2 and RPA3 patients; Fibroblasts (Fibros), negative control. **C**) Teratoma assay showing the differentiation of RPA1, RPA2 and RPA3 iPSCs into: ectoderm, as determined by the presence of neural tubes (row 1); mesoderm, as determined by the presence of cartilage (row 2); endoderm, as determined by the presence of intestinal epithelium with typical mucous cells (row 3). Scale bar = 50 µm (ectoderm and endoderm) and 100 µm (mesoderm). **D**) Karyotype analysis of RPA1 iPSCs with a normal 46, XX karyotype at P16, and RPA2 and RPA3 iPSCs with a normal 46, XY karyotype at P14 and P19, respectively.

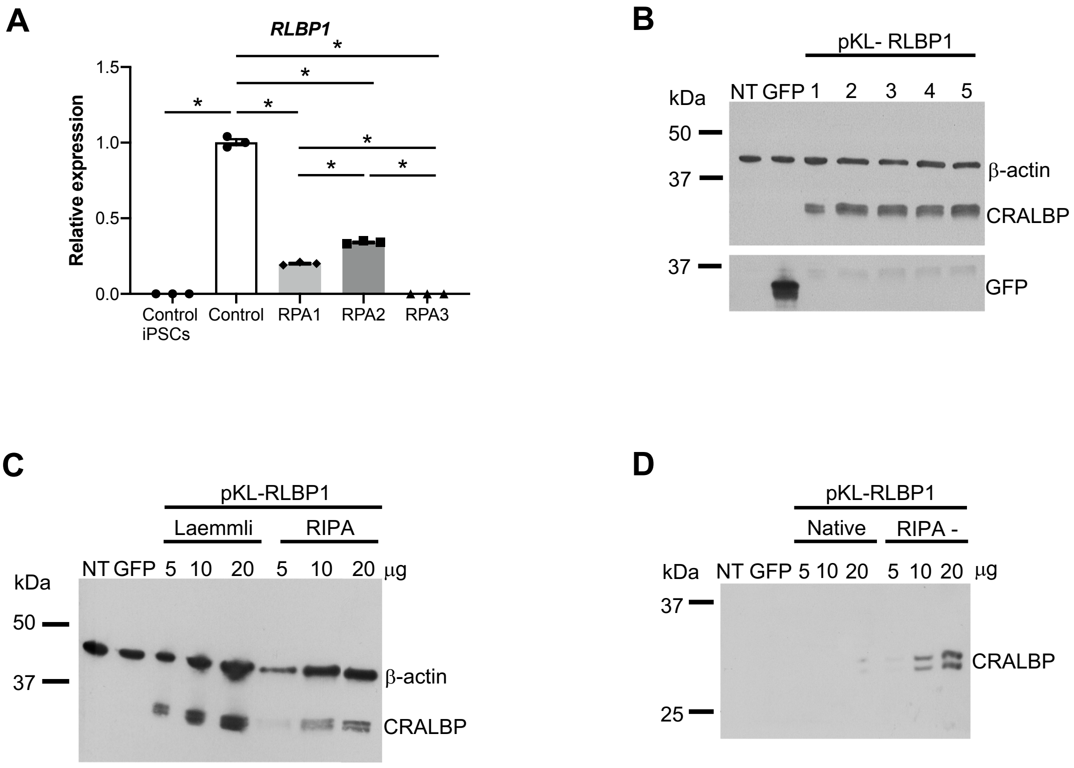


**Figure S2: *RLBP1* in iPSC-derived RPE and CRALBP expression from proviral plasmids.** **A**) qPCR analysis of *RLBP1* expression in control iPSCs, and in control, RPA1, RPA2 and RPA3 iPSC-derived RPE using a F primer spanning the exon 6 to 7 junction and a R primer in exon 7 (situated in the deletion carried by RPA3). Data are represented as mean ± SEM and expressed relative to control; *$p$<0.05; $n$ = 3. **B**) Western blot analysis of CRALBP using a mouse monoclonal antibody in COS-7 cells NT, or transfected with a GFP-expressing plasmid or pKL-RLBP1, and scraped and collected in different combinations of protein inhibitor cocktails (PIC) and buffers: 1. Roche PIC and resuspension in Laemmli buffer (standard conditions); 2. Roche PIC and lysis in RIPA buffer; 3. Cell signalling PIC and lysis in Cell signalling lysis buffer; 4. Cell signalling PIC and lysis in RIPA buffer; 5. Roche PIC and lysis in Cell Signalling buffer; β-actin was used as loading control and GFP as a positive transfection control. **C**) Western blot analysis of CRALBP using a mouse monoclonal antibody in COS-7 cells NT or transfected with a GFP-expressing plasmid or pKL-RLBP1, and collected in Laemmli or RIPA buffer; β-actin was the loading control. **D**) Western blot analysis of CRALBP using a mouse polyclonal antibody in COS-7 cells NT or transfected with a GFP-expressing plasmid or with pKL-RLBP1, collected in Biorad Native sample buffer or lysed in RIPA (without SDS) buffer, and migrated under non-denaturing conditions; β-actin was the loading control.

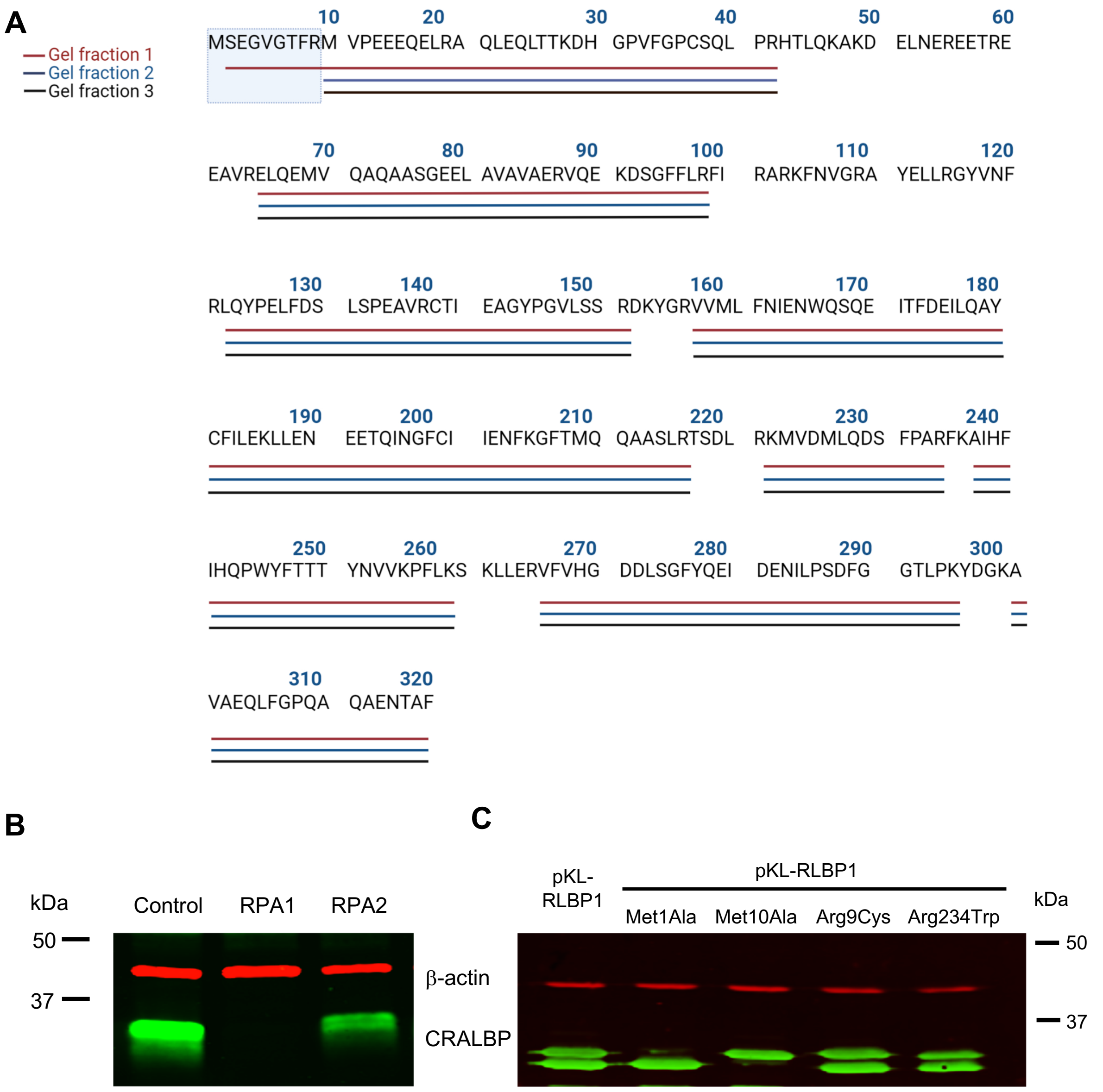


**Figure S3: Proteomic and western blot analysis of control or mutant CRALBP isoforms.** Alignment of the CRALBP peptides identified by mass spectrometry in the different gel fractions on the full length CRALBP sequence. The light blue square indicates the differing N-terminus between the two isoforms. **B**) Western blot analysis of CRALBP in 30 µg lysate of control, RPA1 and RPA2 iPSC-derived RPE cultured in 24-well plates. **C**) Western blot analysis of the CRALBP isoforms in 15 µg lysates of HEK-293 cells transfected with the control pKL-RLBP1 plasmid and the plasmids mutated in the ATG start codons (Met1Ala, Met10Ala) or carrying the missense variants present in patients RPA1 (Arg234Trp) and RPA2 (Arg9Cys); β-actin was the loading control in B and C.

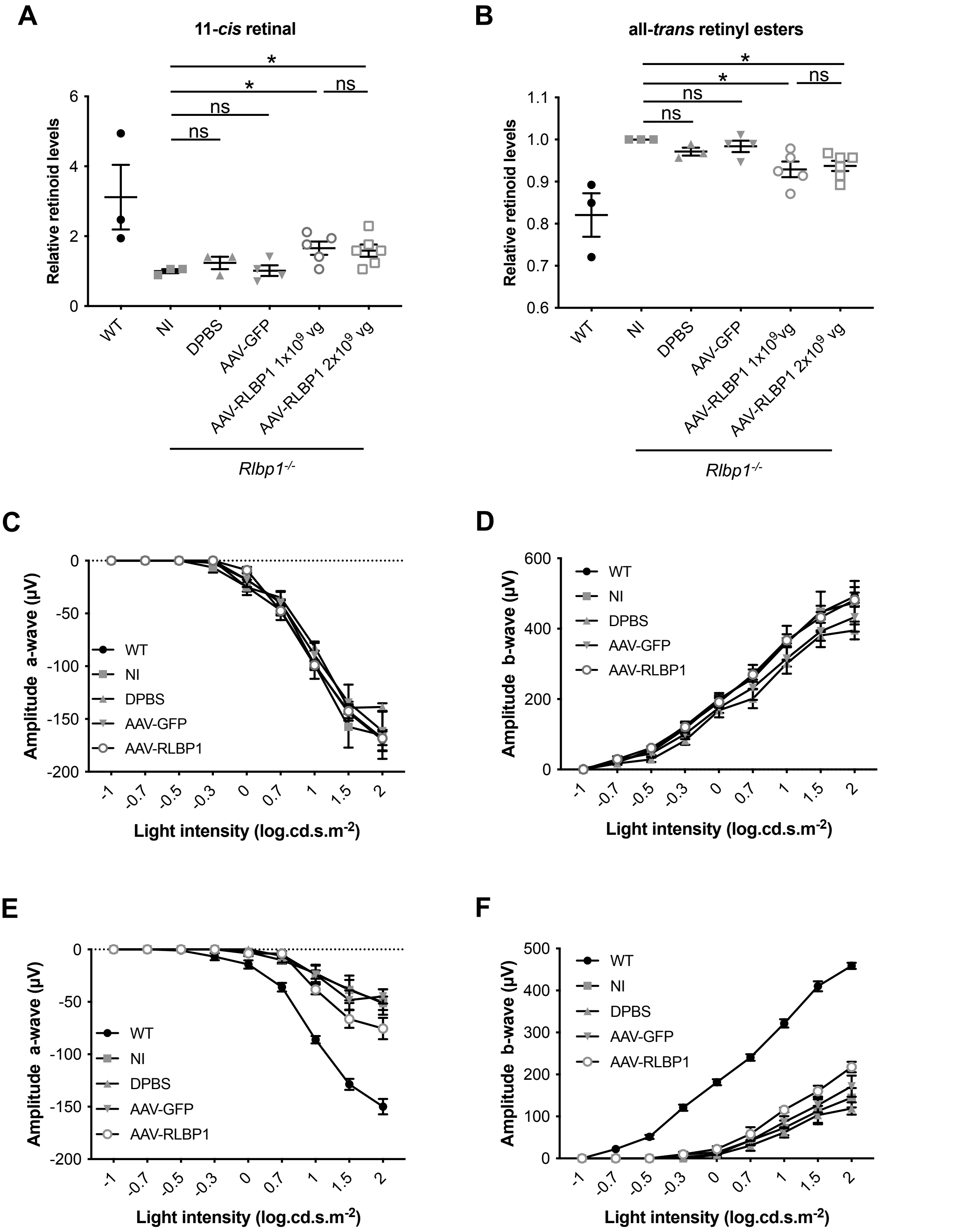


**Figure S4: Analysis of retinoid levels and visual function in *Rlbp1$^{-/-}$* mice.** HPLC of 11-*cis* retinal (**A**) and all-*trans* retinal ester (**B**) levels in wildtype (WT) and *Rlbp1$^{-/-}$* mice that were non-injected (NI) or injected with DPBS, AAV-GFP or 2 doses ($1x10^9$ vg and $2x10^9$ vg) of AAV-RLBP1 at 7 weeks post-treatment. All data are represented as mean ± SEM and expressed relative to NI *Rlbp1$^{-/-}$* mice; *$p<0.05$; ns, non-significant; *n* is indicated by the symbols. Analysis of a-wave (**C**) and b-wave (**D**) ERG amplitudes (in µV) prior to photobleaching (baseline) in WT and NI, DPBS-injected, AAV-GFP-injected and AAV-RLBP1 ($1x10^9$ vg)-injected *Rlbp1*$^{-/-}$ mice at 8 weeks post-treatment following stimulation at increasing light intensities expressed in log candela second/metre$^2$. Data are represented as mean ± SEM; *n* = 6 (WT), 5 (NI), 3 (DPBS), 4 (AAV-GFP) and 5 (AAV-RLBP1). Analysis of a-wave (**E**) and b-wave (**F**) ERG amplitudes (in µV) following photobleaching at increasing light intensities expressed in log candela second/metre$^2$ (log.cd.s.m$^{-2}$) and dark-adaptation of wild type (WT) and *Rlbp1*$^{-/-}$ mice that were NI or injected with DPBS, AAV-GFP or $1x10^9$ vg AAV-RLBP1 at 10 weeks post-treatment. Data are represented as mean ± SEM.

**Table S1. PCR, sequencing, genotyping and mutagenesis primers**

| **Target** | **Experiment** | **Sequence (5' to 3')** |
|---|---|---|
| *Mlu*I-*RLBP1*<br>*RLBP1*-*Xho*I | Vector production | F: **ACG CGT** ATG TCA GAA GGG GTG GGC AC<br>R: **CTC GAG** TCA GAA GGC TGT GTT CTC AG |
| pKL backbone<br>*RLBP1* exon 5<br>*RLBP1* exon 7<br>*RLBP1* exon 5<br>*RLBP1* exon 6<br>RLBP1 exon 7-8 | Sanger sequencing cDNA | F: AAT CTG TGC GGA GCC GAA AT<br>F: CGC GCA CGG AAG TTC AAC GT<br>F: CAA GGG CTT TAC CAT GCA GC<br>R: GAA GCC GCT GTC CTT CTC TT<br>R: AGG CTG TCA AAG AGC TCA GG<br>R: TCC TGG AGC ATG TCC ACC AT |
| *Rlbp1* | Genotyping | F: TTA GAC TCA CAG GGG CCA ACA<br>R1: ATG ATC CTT GGT TGT GAG CTG CTC<br>R2: TAA AGC GCA TGC TCC AGA CT |
| c.1ATG>GCG<br>p.Met1Ala | Site directed mutagenesis | F: TTC GAG CGA ACG CGT **GCG** TCA GAA GGG GTG GG<br>R: CCC ACC CCT TCT GAC GCA CGC GTT CGC TCG AA |
| c.28ATG>GCG<br>p.Met10Ala | Overlapping PCR | F1: TGA TTA ATT CGA GCG AAC GCG T**AT G**TC AGA AGG GGT GGG CAC GTT CCG |
| | | R2: CAG TTG GGC ACG GAG CTC CTG TTC CTC TTC AGG TAC **CGC** GCG GAA CGT GCC CAC C |
| | | F3: GCT CCG TGC CCA ACT GGA GCA GCT CAC AAC CAA GGA CCA TGG ACC TGT CTT TGG CCC |
| | | R4: CTT GGC CTT CTG CAA GGT GTG GCG GGG CAG CTG GCT GCA CGG GCC AAA GAC AGG TCC |
| | | F5: ACC TTG CAG AAG GCC AAG GAT GAG CTG AAC GAG AGA GAG GAG ACC CGG GAG GAG GC |
| | | R6: TGC GCC TGC ACC ATC TCC TGC AGC TCT CGC ACT GCC TCC TCC CGG G |
| | Flanking PCR | F: TGA TTA ATT CGA GCG AAC GCG<br>R: TGC GCC TGC ACC ATC |
| c.25C>T<br>p.Arg9Cys | Site directed mutagenesis | F: GGG TGG GCA CGT TCT GCA TGG TAC CTG AAR'<br>R: 5'TTC AGG TAC CAT GCA GAA CGT GCC CAC CC3 |
| c.700C>T<br>p.Arg234Trp | Site directed mutagenesis | F: GAT TCC TTC CCA GCC TGG TTC AAA GCC ATCC<br>R: GGA TGG CTT TGA ACC AGG CTG GGA AGG AAT C |
| c.25C>T in exon 4<br>(p.Arg9Cys; RPA2) | Sanger sequencing gDNA | F: GAC CCC ACA AAA GGA GGA GG<br>R: GCT GGA CCC TTT TCA CAG GA |
| c.333T>G in exon 5<br>(p.Tyr111X; RPA1 and RPA2) | Sanger sequencing gDNA | F: CCT CAC CCG CAC CTA AGT TT<br>R: GGG GGT CTG GAG GGG AAA TT |
| c.700C>T in exon 8<br>(p.Arg234Trp; RPA1) | Sanger sequencing gDNA | F: TTG CTG GCC TGG AAA TAG GA<br>R: GGT GCC CTA AGG ATG AGG GT |
| Exons 7-9del (RPA3) | Long-range PCR (gDNA) | F: TGT GAA GCT GAG CAC GTC AGA T<br>R: TTG GGA GAA CTT TGG CAT G |

F – forward, R – reverse

**Table S2. RT-PCR and qPCR primer sequences**

| Target | Experiment | Forward primer | Reverse primer | Efficacity |
|---|---|---|---|---|
| *RLBP1* | qPCR exons 6-7 | GAA ATC ACC TTT GAT GAG AT | TCT TCC TGA GAT CTG AAG TC | 2.18 |
| *RLBP1* | qPCR exon 6 | ACC CTG AGC TCT TTG ACA GC | TGA AGA GCA TGA CCA CTC GG | 2.02 |
| SeV | RT-PCR | GGA TCA CTA GGT GAT ATC GAG C | ACC AGA CAA GAG TTT AAG AGA TAT GTA TC | N/A |
| *KOS* | RT-PCR | ATG CAC CGC TAC GAC GTG AGC GC | ACCTTGACAATCCTGATGTGG | N/A |
| *KLF4* | RT-PCR | TTC CTG CAT GCC AGA GGA GCC C | AAT GTA TCG AAG GTGCTC AA | N/A |
| *c-MYC* | RT-PCR | TAA CTG ACT AGC AGG CTT GTC G | TCC ACA TAC AGT CCT GGA TGA TGA TG | N/A |
| *NANOG* | qPCR | CAA AGG CAA ACA ACC CAC TT | TCT GCT GGA GGC TGA GGT AT | 2.06 |
| *OCT3/4* | qPCR | GTA CTC CTC GGT CCC TTT CC | CAA AAA CCC TGG CAC AAA CT | 1.94 |
| *LIN28A* | qPCR | GGG GAA TCA CCC TAC AAC CT | CTT GGC TCC ATG AAT CTG GT | 2.16 |
| *GAPDH* | qPCR | AAC CAT GAG AAG TAT GAC AAC | CTT CCA CGA TAC CAA AGT T | 2.02 |
| *L27* | qPCR | ACG CAA AGC CGT CAT CGT GAA G | CTT GGC GAT CTT CTT CTT GCC | 2.09 |

N/A – non-applicable

**Table S3: Human primary and secondary antibodies**

| Primary antibodies | Host | Clonality | Dilution | Company | Cat # |
|---|---|---|---|---|---|
| anti-ARL13B | Rabbit | Polyclonal | 1/3000 | Proteintech | 17711-1-AP |
| anti-β-actin, clone AC-74 | Mouse | Monoclonal | 1/10000 | Sigma-Aldrich | A5316 |
| anti-BEST1 | Mouse | Monoclonal | 1/500 | Abcam | Ab2182 |
| anti-CRALBP, clone B2 | Mouse | Monoclonal | 1/1000 | Abcam | ab15051 |
| anti-CRALBP | Rabbit | Monoclonal | 1/1000 | Abcam | ab183728 |
| anti-CRALBP | Mouse | Polyclonal | 1/1000 | AgroBio | Directed against recombinant human CRALBP |
| anti-GFP | Rabbit | Polyclonal | 1/2000 | Invitrogen | A6455 |
| anti-LRAT | Rabbit | Polyclonal | 1/250 | Abcam | ab166784 |
| anti-MERTK, clone Y323 | Rabbit | Monoclonal | 1/250 | Abcam | ab52968 |
| anti-Perilipin-2 | Guinea pig | Polyclonal | 1/2000 | Progen | GP47 |
| anti-RPE65 | Mouse | Monoclonal | 1/1000 | Abcam | ab13826 |
| anti-ZO-1 | Rabbit | Polyclonal | 1/100 | ThermoFisher Scientific | 40-2200 |

| Secondary antibodies | Host | Dilution | Company | Cat # |
|---|---|---|---|---|
| anti-Mouse IgG-HRP | Sheep | 1/10000 (WB) | Jackson ImmunoResearch | 515-035-003 |
| anti-Guinea Pig-HRP | Goat | 1/10000 (WB) | Jackson ImmunoResearch | 106-035-003 |
| anti-Mouse IgG IRDye 680RD | Donkey | 1/20000 (WB) | LI-COR Biosciences | 926-68072 |
| anti-Rabbit IgG IRDye 800CW | Donkey | 1/20000 (WB) | LI-COR Biosciences | 926-32213 |
| anti-Mouse IgG Alexa Fluor 488 | Donkey | 1/500 | ThermoFisher Scientific | A-21202 |
| anti-Mouse IgG Alexa Fluor 594 | Donkey | 1/500 | ThermoFisher Scientific | A-21203 |
| anti-Rabbit IgG Alexa Fluor 488 | Donkey | 1/500 | ThermoFisher Scientific | A-21206 |
| anti-Rabbit IgG Alexa Fluor 594 | Donkey | 1/500 | ThermoFisher Scientific | A-21207 |
| anti-Mouse IgG Alexa Fluor 546 | Goat | 1/500 | ThermoFisher Scientific | A10036 |
| anti-Mouse IgG Alexa Fluor 594 AffiniPure | Donkey | 1/500 | Jackson ImmunoResearch | 715-585-150 |
| anti-Rabbit IgG Alexa Fluor 594 AffiniPure | Donkey | 1/500 | Jackson ImmunoResearch | 711-585-152 |
| anti-Mouse IgG Alexa Fluor 488 AffiniPure | Donkey | 1/500 | Jackson ImmunoResearch | 715-546-151 |
| anti-Rabbit IgG Alexa Fluor 488 AffiniPure | Donkey | 1/500 | Jackson ImmunoResearch | 711-545-152 |
